# The radiation of nodulated *Chamaecrista* species from the rainforest into more diverse habitats has been accompanied by a reduction in growth form and a shift from fixation threads to symbiosomes

**DOI:** 10.1101/2023.12.20.572614

**Authors:** Patricia Alves Casaes, José Miguel Ferreira dos Santos, Verônica Cordeiro Silva, Mariana Ferreira Kruschewsky Rhem, Matheus Martins Teixeira Cota, Sergio Miana de Faria, Juliana Gastaldello Rando, Euan K. James, Eduardo Gross

## Abstract

All non-mimosoid nodulated genera in the legume subfamily Caesalpinioideae confine their rhizobial symbionts within cell wall-bound “fixation threads” (FTs). The exception is the large genus *Chamaecrista* in which shrubs and subshrubs house their rhizobial bacteroids more intimately within symbiosomes, whereas large trees have FTs. This study aimed to unravel the evolutionary relationships between *Chamaecrista* growth habit, habitat, nodule bacteroid type, and rhizobial genotype. The growth habit, bacteroid anatomy, and rhizobial symbionts of 30 nodulated *Chamaecrista* species native to different biomes in the Brazilian state of Bahia, a major centre of diversity for the genus, was plotted onto an ITS-*TrnL-F-* derived phylogeny of *Chamaecrista*. The bacteroids from most of the *Chamaecrista* species examined were enclosed in symbiosomes (SYM-type nodules), but those in arborescent species in the section *Apoucouita*, at the base of the genus, were enclosed in cell wall material containing homogalacturonan (HG) and cellulose (FT-type nodules). Most symbionts were *Bradyrhizobium* genotypes grouped according to the growth habits of their hosts, but the tree, *C. eitenorum,* was nodulated by *Paraburkholderia*. *Chamaecrista* has a range of growth habits that allow it to occupy several different biomes and to co-evolve with a wide range of (mainly) bradyrhizobial symbionts. FTs represent a less intimate symbiosis linked with nodulation losses, so the evolution of SYM-type nodules by most *Chamaecrista* species may have (a) aided the genus-wide retention of nodulation, and (b) assisted in its rapid speciation and radiation out of the rainforest into more diverse and challenging habitats.

## Introduction

The monophyletic legume genus *Chamaecrista* (L.) Moench (Leguminosae - Caesalpinioideae) has its centre of diversification in South America (Conceição et al., 2009). Most species occur in Brazil, where 268 of approximately 366 recognized species are distributed in various biomes and vegetation types. About 223 species are considered as Brazilian endemics (LPWG 2020; Rando et al., 2020) particularly in the states of Bahia (BA) and Minas Gerais (MG) which harbour 94 species distributed in such diverse environments as savannah (Cerrado), Campo rupestre (upland rocky fields), semiarid ecosystems (Caatinga), and tropical rain forest (Coutinho et al., 2016; Rando et al., 2016, 2020).

*Chamaecrista* is the ninth largest genus in the Leguminosae (Fabaceae) and the third largest in the Caesalpinioideae subfamily (after *Acacia* and *Mimosa*); it contains a wide variety of plant growth habits and sizes, ranging from trees through to shrubs, and subshrubs/woody herbaceous perennials (Lewis, 2005; Coutinho et al., 2016; Mendes et al. 2017; LPWG 2021). Furthermore, the distribution of *Chamaecrista* is unique in being the only nodulated caesalpinioid genus which has species that have colonized temperate regions (Sprent et al., 2013). All *Chamaecrista* species so far examined form symbiotic root nodules with nitrogen-fixing bacteria collectively known as rhizobia (Gyaneshwar et al., 2011; Peix et al., 2015; Sprent et al., 2017), while related genera comprising the Cassia clade (LPWG 2017), p. ex. *Cassia* L. and *Senna* Mill. do not nodulate (Sprent, 2001, 2009). The independent rise of nodulation and its variation in *Chamaecrista* (Delaux et al., 2015; Naisbitt et al., 1992) suggests that the genus has a pivotal position in the evolution of nodulation (Sprent et al., 2013), and could be used as a model for detailed studies of interactions between plants and nitrogen-fixing bacteria (Singer et al., 2009). Indeed, it is for this reason that Sprent et al. (2013) suggested studying the nitrogen-fixing nodules across *Chamaecrista* species in more depth with a focus on their structure and rhizobial symbionts.

There is a paucity of information about nodule anatomy and development in the paraphyletic grade comprising the Caesalpinioideae (*i.e.,* excluding the Mimosoid clade), which includes *Chamaecrista*. What we do know is that all nodules so far studied from the nine known nodulating non-Mimosoid Caesalpinioideae genera (*Campsiandra*, *Chamaecrista*, *Dimorphandra*, *Dinizia*, *Erythrophleum*, *Jacqueshuberia*, *Melanoxylon*, *Moldenhawera* and *Tachigali*) are indeterminate, retaining meristematic activity (Sprent, 2009; Fonseca et al., 2012; Sprent et al., 2013, 2017; Faria et al. 2022). Moreover, while the nodules in most papilionoid and all mimosoid species so far examined, have their symbiotic rhizobia (bacteroids) released into membrane-bound vesicles called symbiosomes (Sprent, 2009; Sprent et al., 2013, 2017), some papilionoid, and all nodulated (non-mimosoid) caesalpinioid trees so far studied have their bacteroids confined within cell wall-bound modified infection threads termed ‘‘persistent infection threads’’ (PITs) or ‘‘fixation threads’’ (FTs) (Faria et al., 1987, 2022; Naisbitt et al., 1992; Sprent, 2009; Fonseca et al., 2012; Sprent et al., 2013, 2017). In *Chamaecrista*, however, nodules are anatomically diverse with their ultrastructure being apparently related to plant growth habit i.e. the rhizobial bacteroids are enclosed in FTs in tree species, but within membrane-bound symbiosomes in subshrub/woody herbaceous species (henceforth collectively termed “subshrubs”), while larger shrub/treelet species have nodules with intermediate structures between FTs and symbiosomes (Naisbitt et al., 1992).

The nodule anatomy of most species of *Chamaecrista* is so far undescribed, as is the nodulation status of the majority of the genus. In addition, there have been significant advances in microscopy techniques since Naisbitt et al., (1992), particularly in methods to determine the composition of membranes and cell walls that could be associated with the FTs and symbiosomes (Faria et al. 2022). Therefore, the first aim of this study was to record the nodulation status of a wide variety of *Chamaecrista* species covering their whole range of growth habits, from 20 m-high tree species to small subshrubs at only 20-30 centimeters in height. Plant size variation in *Chamaecrista* is also related to the biomes within which they occur: large trees are distributed in tropical rainforests where soils are nitrogen (N) rich, while shrubs and subshrubs are distributed in tropical savannas and in xeric formations with N-poor soils. Moreover, the phylogenetic relations within the genus *Chamaecrista* also shows that monophyletic groups share similar types of habitats and growth habit. The few arborescent species in the genus, all native to tropical rainforests, are grouped in the basal monophyletic section *Apoucouita* (Coutinho et al., 2016; Souza et al. 2021), while the considerably more numerous species from open areas are shrubs and subshrubs that are scattered widely across the taxonomy of the genus (Conceição et al., 2009, Souza et al., 2021). A second aim was to evaluate the anatomy and ultrastructure of root nodules from *Chamaecrista* species of all growth habits, focusing on the chemical composition of structures (symbiosomes, FTs, and intermediates) enclosing the bacteroids within the infected tissue of the nodules.

The third aim was to explore the identity of the rhizobial symbionts of a wide variety of *Chamaecrista* species covering the whole range of growth habits from trees through to shrubs and subshrubs, and from nodules with FTs to those with symbiosomes. Current data suggest that non-mimosoid Caesalpinioideae are preferentially nodulated by *Bradyrhizobium* strains (Fonseca et al., 2012; Yao et al., 2014, 2015; Parker 2015; Sprent et al., 2017; Rathi et al., 2018; Cabral Michel et al., 2021). Indeed, in the specific case of the largest nodulating genus in this group, *Chamaecrista*, several studies have shown that the North American subshrub *C. fasciculata* (Michx.) Greene (partridge pea) has nodules that are associated with *Bradyrhizobium* (Parker 2012, Parker & Rousteau 2014; Urquiaga et al., 2019; Klepa et al., 2019), and a recent molecular analysis of 47 strains from nine shrub and subshrub *Chamaecrista* species in Brazil described all the symbiotic strains as belonging to genotypes of *Bradyrhizobium* (Santos et al., 2017). Outside the New World, reports on native shrubby *Chamaecrista* species in India and Africa suggest that they are also mainly nodulated by bradyrhizobia (de Lajudie et al., 1998; Beukes et al., 2016; Rathi et al., 2018). Although evidence to date suggests that a strong association between *Chamaecrista* and bradyrhizobia is consistent, nothing is known about the diversity of rhizobial symbionts in nodules of tree species which have their bacteroids enclosed in FTs as opposed to those with symbiosomes, nor whether there is a link between rhizobial genotypes and the different biomes within which their hosts occur.

Using these data plus unpublished anatomical data obtained from *Chamaecrista* nodules sampled during the expedition of Sprent et al. (1996), combined with data from the literature (Naisbitt et al. 1992; Santos et al. 2017; Faria et al. 2022), we then test the hypothesis that the distribution of FTs and symbiosomes in the genus is not random and may be the result of co-evolution between groups (sections) of *Chamaecrista* that are native to particular environments (biomes), and the rhizobial microsymbionts that live within them. This was done by constructing a phylogeny with ITS-*TrnL-F* sequences from 119 separate *Chamaecrista* taxa, and then plotting onto it plant growth habit, nodule ultrastructure (occurrence of FTs or symbiosomes, if known), and the *nodC* genotypes of the rhizobial microsymbionts.

## Materials and methods

### Botanical material

To sample root nodules in the field, we first located the 17 *Chamaecrista* species studied here in different biomes of Bahia State, Brazil (Fig. S1). Samples of the root system bearing nodules were collected from each individual plant, mainly during the rainy season, when nodules are most active (dos Reis Junior et al., 2010). Aerial parts of each species were collected, dried, and deposited in the Herbarium of the Universidade Estadual de Santa Cruz (UESC) (Table S1) for confirmation of their identities.

On average, 12 nodules were collected from each of the 17 *Chamaecrista* species, totaling 204 nodules. These samples were used to characterize the infected tissues and to verify the presence of FTs using scanning electron microscopy (SEM), transmission electron microscopy (TEM), light and fluorescence microscopy (see details below).

### Anatomy, histochemistry, ultrastructure and immunocytochemistry of nodules

The nodules were separated from the root system and cut into 1-2 mm^3^ pieces with a razor blade, before being fixed in a solution of 2.5% glutaraldehyde in 0.1 M sodium cacodylate buffer (pH 7.2). Sample processing followed Santos et al. (2017) for anatomical and ultrastructural characterization.

Histochemistry using the fluorescent compound calcofluor white, which binds to beta 1-3 and beta 1-4 polysaccharides, such as those found in cellulose (Wood et al., 1983), was used to provide evidence for the presence of invasive infection threads (ITs) and FTs within the infected tissue of *Chamaecrista* nodules. Briefly, sections of nodules were incubated with calcofluor white (1 g L^-1^) in Evans blue as a background stain (0.5 g L^-1^) (Sigma-Aldrich) according to Wood et al. (1983), and the sections were then observed under a Leica DM2500 equipped with ebq100-04 fluorescence coupled to a Leica DFC310 Fx digital camera.

For transmission electron microscopy (TEM), serial ultrathin sections were collected on nickel grids for immunogold tests using the monoclonal antibody JIM5 to verify the presence of a homogalacturonan (HG) epitope in FTs inside the nodule infected cells; this HG epitope is an essential component of pectin in cell walls (VandenBosch et al., 1989; Fonseca et al., 2012). For species from which *Paraburkholderia* were isolated as potential symbionts, nodules were also tested for the *in-situ* presence of symbiotic strains using a polyclonal antibody against *P. phymatum* STM815^T^ according to dos Reis Junior et al. (2010).

For scanning electron microscopy (SEM), fixed samples were dehydrated in a graded acetone series to absolute, and completely dried using a Bal-Tec CPD 030 critical point drier. Dried samples were mounted on stubs, coated with gold in a Bal-Tec SCD 050 sputter coater and viewed with a FEI Quanta 250 at the Centro de Microscopia Eletrônica (CME) at UESC.

### Rhizobia isolation, cultivation and characterization

Potential rhizobia were isolated from root nodules of seven of the 17 *Chamaecrista* species sampled in the field from native undisturbed environments. An intensive effort was made to isolate rhizobia from tree species of *Chamaecrista* because they often have woody and lignified nodules, and hence the symbiotic bacteria are more difficult to isolate. Accordingly, approximately 230 nodules were used for rhizobial isolation from *C. ensiformis* var. *plurifoliolata*, *C. duartei*, *C. eitenorum* and *C. bahiae*. Rhizobia were isolated, cultivated and characterized following the procedures of Rhem et al. (2021).

### Bacterial DNA extraction, amplification, sequencing and phylogenetic analysis

For samples from each plant, bacterial isolates were grouped and selected according to similarities in their phenotypical (colony) characteristics. Genomic DNA from selected isolates was extracted according to Santos et al. (2017). The DNA was resuspended in ultrapure water and stored at −20 °C. The yield and purity of the extracted DNA was measured in a spectrophotometer (Shimadzu, SPD-M6A) by the ratio of absorbance at 260 and 280 nm. For *Bradyrhizobium* strains, the 16S rRNA gene and ITS region were amplified. A multilocus sequence analysis (MLSA) was performed following Rhem et al. (2021) to identify the phylogenetic positions of symbiont strains from different *Chamaecrista* species. For MLSA, DNA sequences of four housekeeping genes were used, i.e., *recA* encoding recombinase A, *dnaK* encoding the Hsp70 chaperone, *rpoB* encoding RNA polymerase beta subunit, and *glnII* encoding glutamine synthetase isoform II. We also amplified two symbiotic genes, i.e. approximately 930 bp of the *nodC* (nodulation N-acetylglucosaminyltransferase) and 780 bp of the *nifH* (nitrogenase reductase) genes. Thermal cycler programs and primers used are described in Rhem et al. (2021). For *Paraburkholderia* strains, the 16S rRNA, *recA*, *nifH* and *nodC* genes were amplified with the same primers and PCR conditions cited by Silva et al. (2018). Amplified DNA was verified by horizontal electrophoresis in 1% (w/v) agarose gels and PCR products were purified following a cold salt precipitation and resuspended in sterile ultrapure water. All amplicons obtained were sequenced by ACTGene Análises Moleculares Ltda (Alvorada, RS, Brazil). Nucleotide sequences of strains were analyzed for percentage sequence similarity using BLASTn of the National Center for Biotechnology Information (NCBI). Sequences from the present study and those of closely related, reference and type strains (as per the LPSN list of valid and not validly published type strains) were downloaded from NCBI in FASTA format, and then aligned using CLUSTAL X (Thompson et al. 1997) of MEGA X (Kumar et al. 2018). Phylogenetic trees were constructed with the sequence alignments of all the tested genes. Maximum-likelihood trees were built in MEGA 7 using the Tamura 3-parameter correction method. The robustness of the branches of the trees was estimated with 1000 bootstrap replications. The partial sequences of all genes derived from the present study were deposited in GenBank, and their Accession Numbers are listed in Table S2. Please note that it was not possible to amplify all of the examined gene loci from all of the strains.

### Plant DNA extraction, amplification, sequencing and phylogenetic analysis

DNA was sampled from 106 species of *Chamaecrista* (119 taxa) covering all sections and including all species with root nodules analyzed in the present study plus others accessed from the literature and from the dataset associated with Sprent et al. (1996), with only *C. zygophylloides* (Taub.) H.S. Irwin & Barneby not included (Table S3). Additionally, four species of *Cassia* and *Senna* were included as an outgroup. Most of the sequence data were accessed from GenBank (http://www.ncbi.nlm.nih.gov/genbank/), but seven new sequences of the plastid *trnL-F* and four of the nuclear internal transcribed spacer (ITS) were generated for a further seven *Chamaecrista* species. All DNA sequences and associated voucher information are deposited in GenBank (Table S3). Total DNA extraction, amplification, PCR product purification, and sequencing were as described in Conceição et al. (2009). Electropherograms were assembled and edited using the Geneious platform (Drummond et al., 2012). Alignments of all sequences were performed using Muscle (Edgar, 2004) with default settings. Manual edition to correct obvious alignment errors and to remove sections with dubious alignments were inspected using the Geneious platform (Drummond et al., 2012). Bayesian analyses (BAs) were performed with MrBayes 3.1 (Ronquist & Huelsenbeck, 2003) using a combined data set with four partitions (*trnL-F*, ITS1, 5.8S and ITS2). Nucleotide-substitution models were selected, on the basis of the Akaike information criterion (AIC) values, with JModeltest 2.1 (Guindon & Gascuel, 2003; Darriba et al., 2012). The substitution models selected for *trnL-F* was GTR+G and for ITS1 was GTR+I+G, for 5.8S was SYM+I and for ITS2 was GTR+G. Indels were coded as the standard characters “variable”. We run all phylogenetic analyses via the CIPRES Science Gateway v. 3.3 online portal (Miller et al., 2010). We used FigTree 1.4.2 (Rambaut, 2014) to view and edit the final tree.

### Nodulation tests with rhizobial strains

We selected strains isolated from *Chamaecrista* species to evaluate their nodulation capacity. Six *Chamaecrista* species, ranging in size/habit from trees (*C. bahiae, C. ensiformis* var. *plurifoliolata* and *C. duartei*), treelets (*C. blanchetii*) to subshrubs (*C. desvauxi, C. rotundifolia*), as well as the promiscuous papilionoid legume Siratro (*Macroptilium atropurpureum*) were used as host plants in the nodulation tests. In addition, *Mimosa pudica* was used as a promiscuous mimosoid legume to test the two *Paraburkholderia* isolates. Siratro seedlings were inoculated with the different strains according to Santos et al. (2017), while the *Chamaecrista* spp. and *M. pudica* were inoculated according to Silva et al. (2018). At harvest (3 months after inoculation), nodulation of the root system was evaluated for each plant, and the presence of the symbiosis-essential protein leghemoglobin (Lb), as indicated by a pinkish coloration in their interior, was scored.

## Results

### Anatomy, ultrastructure and immunocytochemistry of nodules

*Chamaecrista* species have a wide range of growth habits and habitats, but most are shrubs and subshrubs. The few tree species are usually found in rainforests, such as the *Mata Atlântica* (Fig. 1A) and Amazon Forest; an exception is *C. eitenorum* which has its habitat in seasonally dry tropical forests (SDTF) in the Chapada Diamantina. Most of the tree species have characteristic ramiflorous racemes (Fig. 1B) and are placed in the small section *Apoucouita* at the base of the genus (Coutinho et al., 2016). Woody shrub species occur in the *Cerrado* (savannah) and in open rocky fields often at altitudes >1000 m (denoted *Campo Rupestre*) (Fig. 1C, D); these belong in sect. *Absus*, members of which have terminal or axillary racemes. The *Cerrado* and *Caatinga* (a semiarid Brazilian biome) are also the habitat for smaller, but still woody, sub-shrub and herbaceous species (Fig. 1E, F) which have axillary and supra-axillary reduced racemes (fascicles); these are mostly contained in sect. *Chamaecrista*.

**Fig. 1.**
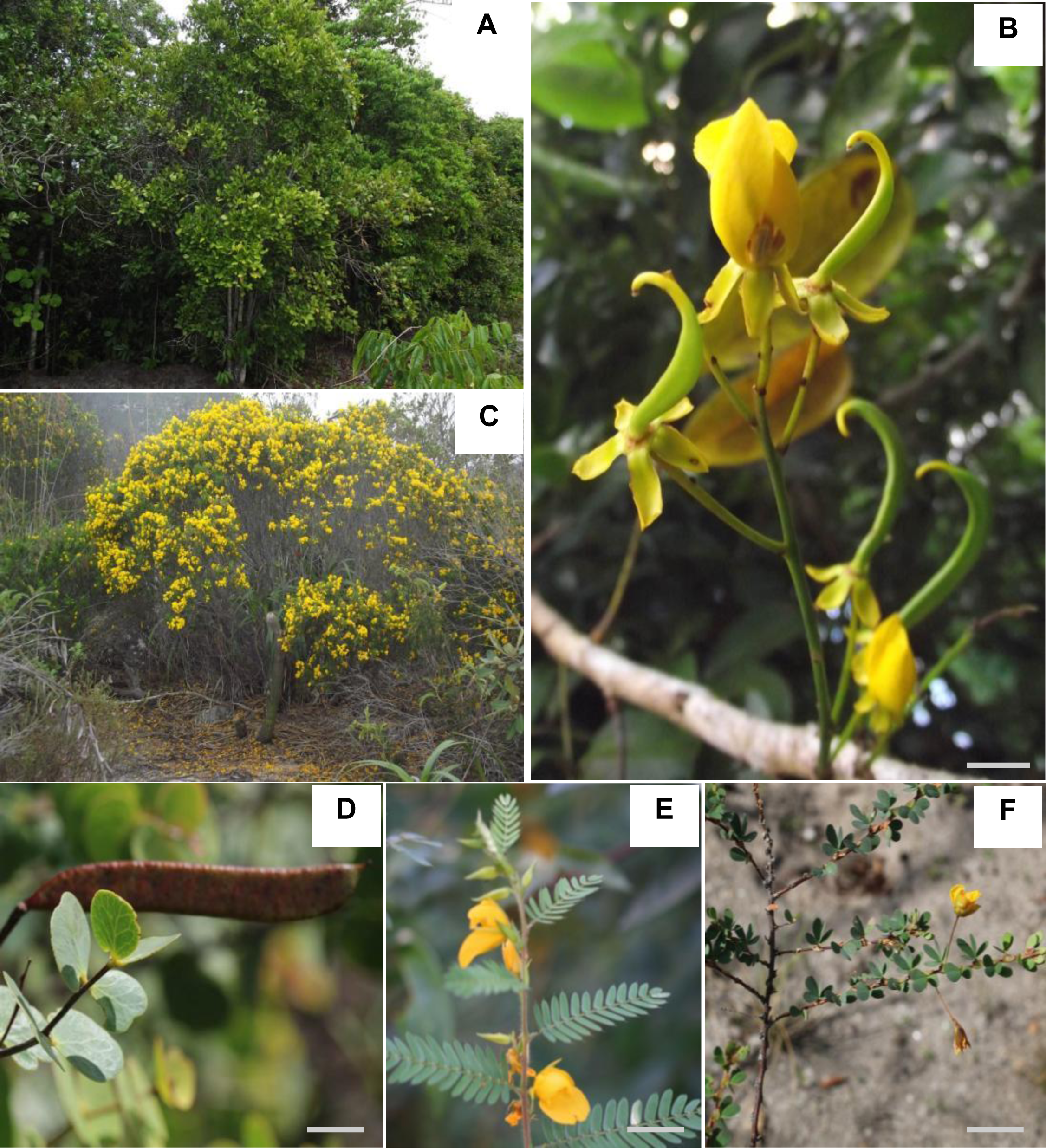
*Chamaecrista* species and their morphological variation across the various sections of the genus. (A) Tree species of *C. bahiae* (Section *Apoucouita*) in the tropical rainforest; (B) Flower and initial developing fruit of *C. duartei*; (C) General view of a *C. confertiformis* plant in *campo rupestre* habitat; (D) Detail of a *C. blanchetii* (Section *Baseophyllum*) fruit; (E) Supra-axillary fascicles of *C. repens* (Section *Chamaecrista*); (F) Solitary flowers of *C. ramose* (Section *Chamaecrista*). Scale bars: (B) = 3 cm; (D, E, F) = 1 cm.

A complete list of nodulating *Chamaecrista* species is given in Table S4. Nineteen *Chamaecrista* taxa were examined in the present study for nodulation, plus twelve from the study of Sprent et al. (1996), and a further eight from de Faria (unpublished). All were nodulated, including 26 new reports.

The nodules sampled in the present study (varying from 0.1 to 2 cm in length) were usually found on the secondary superficial roots (Fig 2A) of all the *Chamaecrista* species examined. In general, nodules were cylindrical when young, but became lobed with age with a few branches, and were a dark brown surface color (Fig. 2B); most were viable and active as evidenced by the pink color in their interior due to Lb production (inset Fig. 2B). Nodule morphology and anatomy characterized all of the samples as indeterminate as they have a persistent meristem at the distal end (Fig. 2C). *Chamaecrista* nodules have a central infected tissue containing the microsymbionts which is surrounded by an uninfected cortex consisting of a parenchymatous outer cortex composed of several layers (4 to 6) of isodiametric cells with phenolic compounds being found scattered throughout the outer cortical region; this is separated from the inner cortex by an endodermis (Fig. 2C). Several vascular bundles are located at the periphery of the inner cortex (Fig. 2C). In all *Chamaecrista* species, the central region of nodules contains a combination of large infected parenchymatic cells and smaller uninfected interstitial cells (Fig. 2C). In the invasion zone, which was located distally to the infected tissue, the rhizobia were contained within ITs that invaded cells (Fig. 2D). Depending on the growth habit of the species, these ITs either released the bacteria into symbiosomes (Fig. 2G, S2), or the ITs developed into thinner-walled FTs which occupied almost the entire cytoplasmic volume of the infected cells (Fig. 2D, S2). In mature nodules, senescent cells were observed in the proximal part of the infected tissue, with a collapsing mass of FTs in their interior (Fig. 2E). The small uninfected interstitial cells were vacuolated, and distinct from the infected cells (Fig. 2F, G, S2). In subshrub *Chamaecrista* species, the root nodules had infected cells with symbiosomes completely occupying the available cytoplasm (Fig. 2G, S2I, K).

**Fig. 2.**
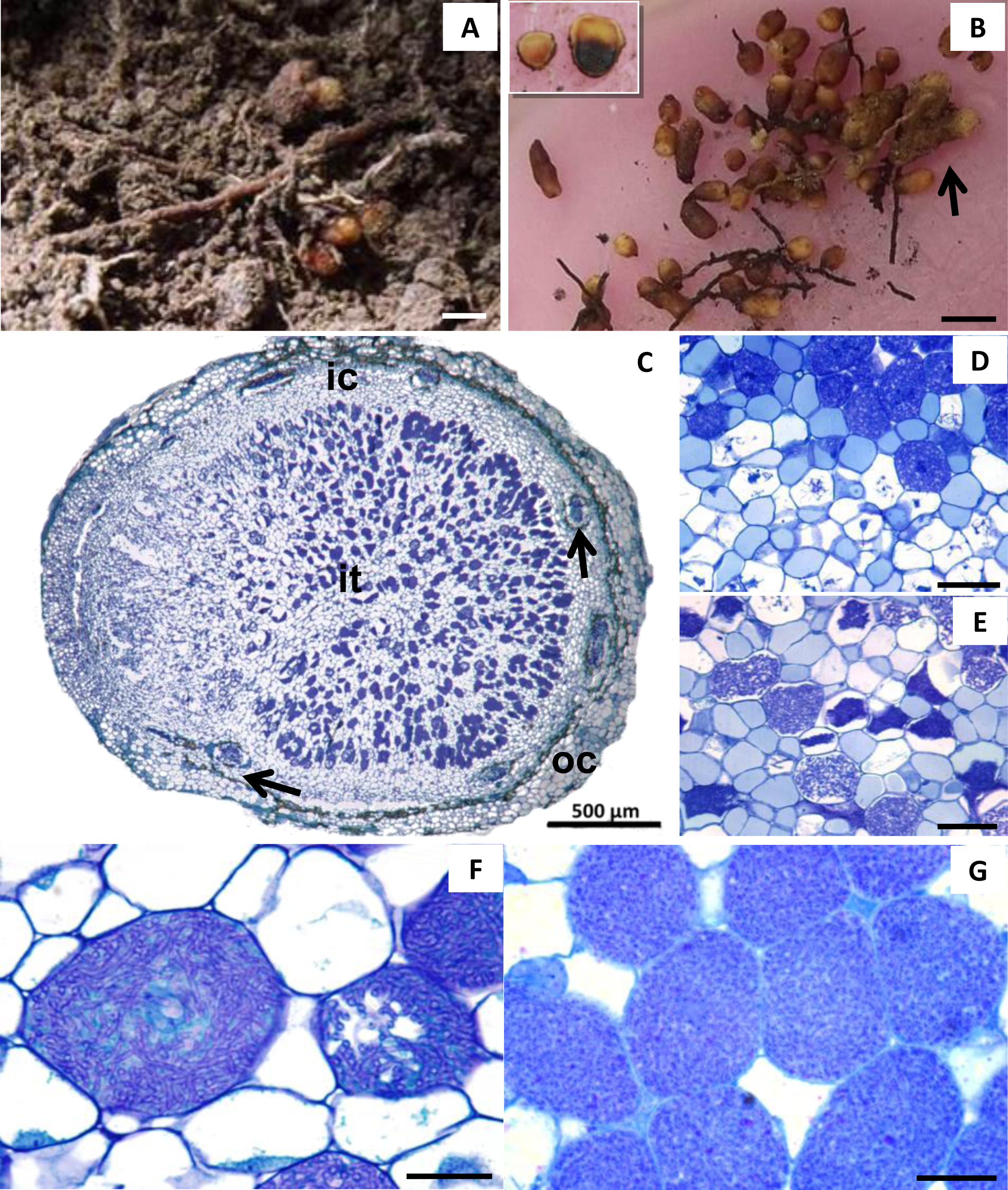
Morphology and anatomy of nodules from species in the various sections of the genus *Chamaecrista*. **A.** Nodules of *C. bahiae* (Section *Apoucouita*) under natural soil conditions; **B.** View of nodule morphologies. An arrow indicates a branched nodule; **C.** General view of a longitudinal section of a *C. bahiae* nodule showing infected tissue (it) in the center surrounded by the inner (ic) and outer cortex (oc); **D.** Sector of a *C. bahiae* nodule showing the invasion zone and part of the nitrogen (N) fixation zone; **E.** Proximal part of a *C. bahiae* nodule showing infected cells in different stages of senescence; **F.** Detail of mature infected cells of *C. bahiae* occupied by fixation threads and interstitial uninfected cells; **G.** Detail of mature infected cells of *C. rotundifolia* (Section *Chamaecrista*) occupied by symbiosomes and some interstitial non infected cells. Scale bars: **A, B** = 1 cm; **C** = 500 μm; **D, E** = 50 μm; **F, G** = 20 μm.

The occurrence of nodules with conspicuous FTs was confined to tree *Chamaecrista* species in the section *Apoucouita* (Table 1, S4, Figs. 3, 4, S2), which are generally restricted to tropical forests (Coutinho et al. 2016; Souza et al. 2021). The composition of the FT cell walls was examined using a combination of three methods: (1) histochemical staining of cellulose using Calcofluor white, (2) anatomical observations with light microscopy and electron microscopy (TEM and SEM), and (3) immunogold labelling of pectin HG using JIM5 combined with TEM.

**Fig. 3.**
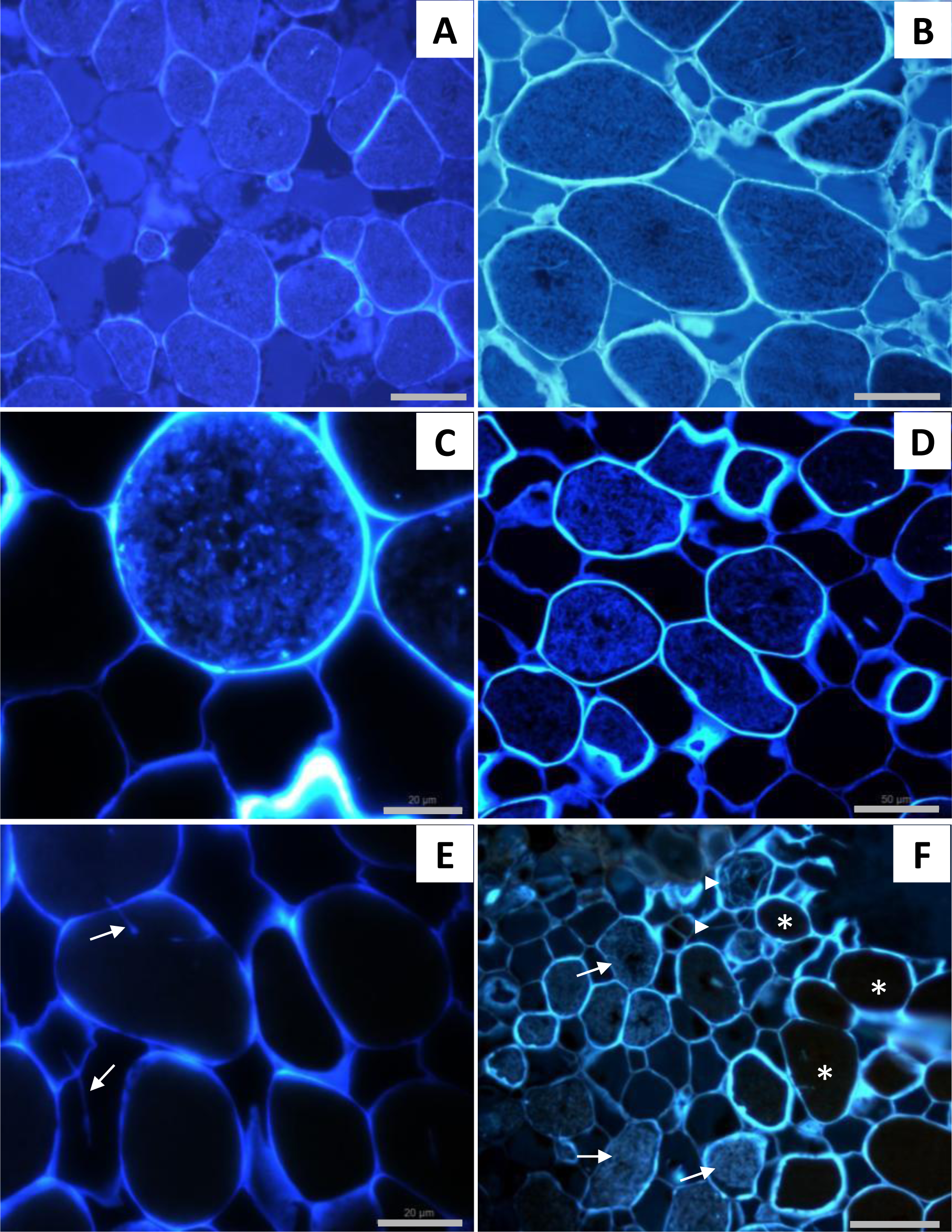
Fluorescence micrographs of Calcofluor White-stained semi-thin sections of nodules from species in the various sections of the genus *Chamaecrista*. Fluorescence was detected in the infected cells of: **A.** *C. bahiae* (Section *Apoucouita*); **B.** *C. duartei* (Section *Apoucouita*); **C.** *C. confertiformis* (Section *Baseophyllum*) and **D.** *C. blanchetiformis* (Section *Baseophyllum*). **E.** No fluorescence was detectable in the infected cells of *C. rotundifolia* (Section *Chamaecrista*). **F.** Fluorescence was detected in the newly-invaded cells of the invasion zone and in the early infected cells, but not in the mature infected cells of *C. belemii* (Section *Absus* subs. *Zygophyllum*). Scale bars: **A, B, D, F** = 50 μm; **C, E**= 20 μm.

**Fig. 4.**
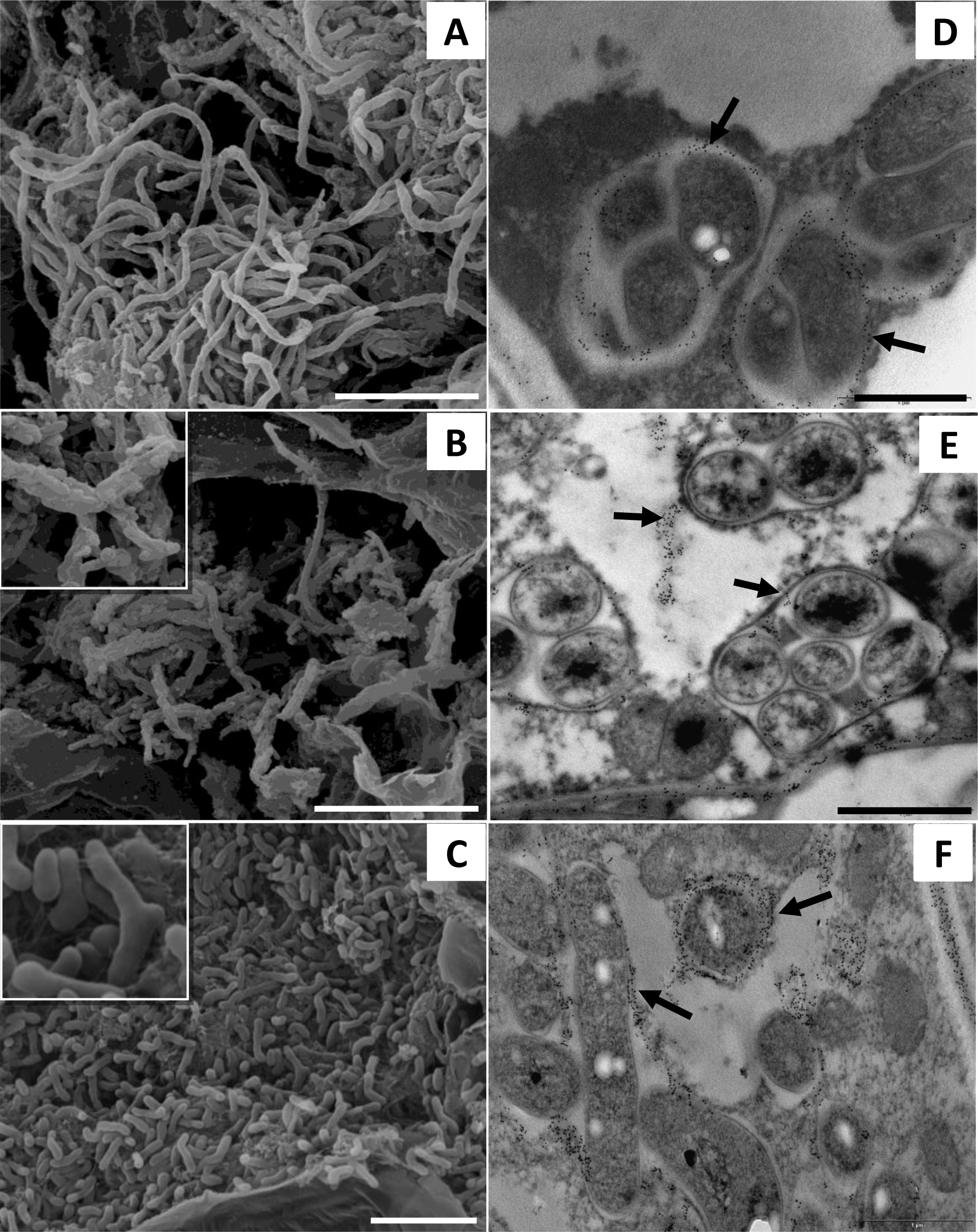
Scanning electron micrographs (SEMs) (**A – C**) and Transmission electron micrographs (TEMs) (**D – F**) of infected cells in *Chamaecrista* nodules from species in the various sections of the genus demonstrating the structure and morphology of the bacteroids across the genus. **A.** *C. bahiae* (FT-type, Section *Apoucouita*); **B.** *C. repens* (FT-SYM-type, Section *Chamaecrista*); **C.** *C. rotundifolia* (SYM-type, Section *Chamaecrista*). **D – F.** TEMs of infected cells after immunogold localization of homogalacturonan epitopes with the monoclonal antibody JIM5, which recognizes a pectin epitope in plant cell walls. **D** *C. bahiae*; **E** *C. arrojadoana* (FT-SYM-type, Section *Chamaecrista*) showing gold particles in symbiosome membrane (wall) and matrix (arrows); **F** *C. supplex* (SYM-type, Section *Chamaecrista*) Scale bars: **A, C** = 10 μm; **E** = 10 μm; **B, D, F**= 1 μm.

**Table 1.**
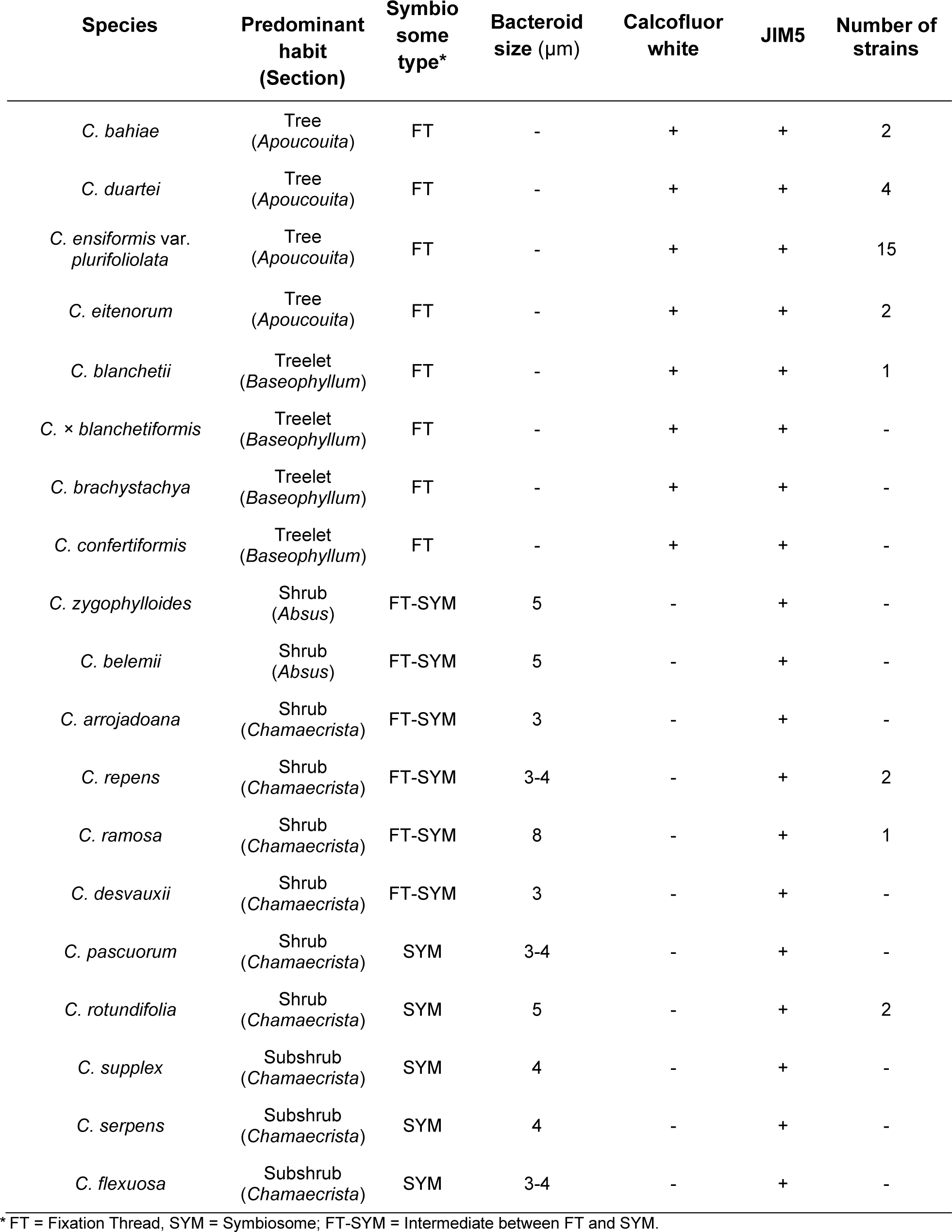
Plant growth habit (and taxonomic section within *Chamaecrista*), plus symbiosome type (fixation threads, symbiosomes or intermediate), bacteroid size, reaction to calcofluor white and JIM5 observed in infected cells of root nodules from *Chamaecrista* spp. native to Bahia State, Brazil, and number of rhizobia strains isolated from each sample.

| Species | Predominant habit (Section) | Symbiosome type* | Bacteroid size (µm) | Calcofluor white | JIM5 | Number of strains |
| --- | --- | --- | --- | --- | --- | --- |
| *C. bahiae* | Tree (Apoucouita) | FT | - | + | + | 2 |
| *C. duartei* | Tree (Apoucouita) | FT | - | + | + | 4 |
| *C. ensiformis* var. *plurifoliolata* | Tree (Apoucouita) | FT | - | + | + | 15 |
| *C. eitenorum* | Tree (Apoucouita) | FT | - | + | + | 2 |
| *C. blanchetii* | Treelet (Baseophyllum) | FT | - | + | + | 1 |
| *C. × blanchetiformis* | Treelet (Baseophyllum) | FT | - | + | + | - |
| *C. brachystachya* | Treelet (Baseophyllum) | FT | - | + | + | - |
| *C. confertifomis* | Treelet (Baseophyllum) | FT | - | + | + | - |
| *C. zygophylloides* | Shrub (Absus) | FT-SYM | 5 | - | + | - |
| *C. belemii* | Shrub (Absus) | FT-SYM | 5 | - | + | - |
| *C. arrojadoana* | Shrub (Chamaecrista) | FT-SYM | 3 | - | + | - |
| *C. repens* | Shrub (Chamaecrista) | FT-SYM | 3-4 | - | + | 2 |
| *C. ramosa* | Shrub (Chamaecrista) | FT-SYM | 8 | - | + | 1 |
| *C. desvauxii* | Shrub (Chamaecrista) | FT-SYM | 3 | - | + | - |
| *C. pascuorum* | Shrub (Chamaecrista) | SYM | 3-4 | - | + | - |
| *C. rotundifolia* | Shrub (Chamaecrista) | SYM | 5 | - | + | 2 |
| *C. supplex* | Subshrub (Chamaecrista) | SYM | 4 | - | + | - |
| *C. serpens* | Subshrub (Chamaecrista) | SYM | 4 | - | + | - |
| *C. flexuosa* | Subshrub (Chamaecrista) | SYM | 3-4 | - | + | - |

\* FT = Fixation Thread, SYM = Symbiosome; FT-SYM = Intermediate between FT and SYM.

Analysis of cellulose deposition in nodule infected cells using calcofluor white showed a similar pattern to the pectin observations *i.e.* with primary cell wall components evident in nodules from trees, treelets and large shrubs, but not in mature infected cells in nodules of subshrubs (Table 2). For example, cellulosic material was detected in FTs in nodules of *C. bahiae* (Fig. 3A), *C. duartei* (Fig. 3B), *C. ensiformis* var. *plurifoliolata*, *C. brachystachya, C. confertifomis* (Fig. 3C) and *C.* x *blanchetiformis* (Fig. 3D). Fluorescence indicating cellulose was also observed in the infected zone of a *C. arrojadoana* nodule (Fig. 3E) where it was localized to an IT in cells recently invaded by rhizobia and, more generally, in immature invaded cells, but not in mature infected cells of this subshrub species (Fig. 3F).

**Table 2.** Nodulation ability on various plant hosts of rhizobial strains isolated from different *Chamaecrista* host species native to Bahia state, Brazil. All strains tested were *Bradyrhizobium* except for those indicated as *Paraburkholderia\**. The *nodC* group that the *Bradyrhizobium* strains belonged to (Fig. 7) are indicated in parentheses: Subshrub-Shrub Cluster I (SSCI), Subshrub-Shrub Clus<u>ter II (SSCII), Tree Cluster I (TCI), Tree Cluster II (TCII), or not known.</u>

| Strain | Host | *Chamaecrista* spp. tested* | Siratro nodulation |
| --- | --- | --- | --- |
| BRUESC623 (SSCII) | *C. blanchetii* (Treelet) | *C. blanchetii* (+) | (-) |
| BRUESC956 (TCII) | *C. ensiformis* var. *plurifoliolata* (Tree) | *C. rotundifolia* (+) | (+) |
| BRUESC1010 (TCI) | *C. ensiformis* var. *plurifoliolata* (Tree) | *C. rotundifolia* (+)<br>*C. ensiformis* (+) | nt |
| BRUESC964 (TCII) | *C. ensiformis* var. *plurifoliolata* (Tree) | *C. ensiformis* (+) | (-) |
| BRUESC967 (TCI) | *C. ensiformis* var. *plurifoliolata* (Tree) | *C. rotundifolia* (+) | (+) |
| BRUESC1011 (not known) | *C. ensiformis* var. *plurifoliolata* (Tree) | *C. bahiae* (-)<br>*C. rotundifolia* (+)<br>*C. ensiformis* (+) | (+) |
| BRUESC952 (not known) | *C. bahiae* (Tree) | *C. rotundifolia* (+)<br>*C. blanchetii* (-)<br>*C. desvauxii* (-) | (+) |
| BRUESC1033 (not known) | *C. repens* (Shrub) | *C. rotundifolia* (+)<br>*C. blanchetii* (-) | (-) |
| BRUESC1034 (singleton) | *C. repens* (Shrub) | *C. bahiae* (+)<br>*C. rotundifolia* (+) | (-) |
| BRUESC1106 (not known) | *C. duartei* (Tree) | *C. blanchetii* (-) | (-) |
| BRUESC1107 (not known) | *C. duartei* (Tree) | *C. desvauxii* (+) | (+) |
| BRUESC1102 (SSCI) | *C. rotundifolia* (Subshrub) | *C. desvauxii* (+) | nt |
| BRUESC1103 (SSCI) | *C. rotundifolia* (Subshrub) | *C. bahiae* (-)<br>*C. blanchetii* (-) | (+) |
| *BRUESC1092 (*Paraburkholderia*) | *C. eitenorum* (Tree) | *C. duartei* (+)<br>*C. desvauxii* (-)<br>*C. ensiformis* (+) | (-) |
| *BRUESC1093 (*Paraburkholderia*) | *C. eitenorum* (Tree) | *C. duartei* (+)<br>*C. desvauxii* (-)<br>*C. ensiformis* (+) | (-) |

nt = not tested; ns = not sequenced. (+), (-) = positive and negative nodulation. \*Also nodulated *Mimosa pudica*. BRUESC955 (TCI), BRUESC961 (TCI), BRUESC963 (TCII), BRUESC965 (TCII), BRUESC966 (not known), BRUESC968 (TCII), BRUESC969 (TCII), and BRUESC1091 (TCI) nodulated Siratro, but were not tested on any *Chamaecrista* species.

Under the SEM, conspicuous FTs were observed in nodules of the trees *C. bahiae* (Fig. 4A), *C. duartei*, *C. eitenorum* and *C. ensiformis* var. *plurifoliolata*, and in the shrub species *C. blanchetiformis*, *C. brachystachya* and *C. confertiformis*, all of which had FTs with well-defined walls like those of the “standard” ITs of many legume types. The mature infected cells in the infected zone of the nodules were densely occupied by FTs, and these differed in structure from those observed in nodules of the subshrub species *C. repens* (Fig. 4B), *C. arrojadoana* and *C. ramosa*, wherein bacteroids were enclosed in FTs with thinner cell walls (so-called FT-SYM; Table S4) which allowed for the bacterial outline with associated material on the bacterial surface to be observed under the SEM (inset Fig. 4B). For nodules on the subshrub species, *C. belemi, C. blanchetii, C. zygophylloides, C. desvauxii*, *C. pascuorum, C. rotundifolia* (Fig. 4C, Table S4) and *C. supplex*, the bacteroids were free in the cells, although they are most likely to be enclosed within symbiosome membranes, most of which did not survive processing for SEM. The SEM observations were confirmed using light microscopy and TEM *i.e*. that tree species in the section *Apoucouita*, such as *C. bahiae*, *C. duartei* and *C. ensiformis* have FTs (Fig. 4D, S2A - D), that symbiosomes in treelets from the section *Baseophyllum* have an intermediate (FT-SYM) structure and/or contain both FTs and symbiosomes (Fig. S2E – H), while shrubs and subshrubs from the sections *Absus* and *Chamaecrista* (Fig. 4E, F, S2I – L) generally contain symbiosomes only. However, there are several exceptions in sections *Absus* and *Chamaecrista* that have an FT-SYM type, such as C. *arrojadoana* (Fig. 4F), *C. mucronata* (Fig. S2L), and *C. desvauxii* (Naisbitt et al. 1992).

Immunocytochemistry using the monoclonal antibody JIM5 provided further information about the nature of FTs and symbiosomes in *Chamaecrista* nodules. It indicated the presence of an unesterified HG epitope in the pectic component of both FTs (Fig. 4D, S2B, D) and symbiosomes (Fig. 4E, S2H, L) in infected cells of nodules from several species of *Chamaecrista* examined (Table 2; Fig. S2). For nodules on tree species in the section *Apoucouita*, such as *C. bahiae*, *C. duartei* and *C. ensiformis*, JIM5 immunogold labeled the walled FTs (Fig. 4D, S2B, D). JIM5 also labelled the thinner walled FTs of treelets and large shrubs in the section *Baseophyllum*, such *as C. blanchetii* (Fig. S2F), but in nodules on other *Baseophyllum* species, such as *C. cytisoides*, the infected cells contained a combination of JIM5-labelled symbiosomes and FTs, with the FTs being more densely labeled (Fig. S2H). Similarly, JIM5 labeling was observed in nodules of some subshrub species in the highly speciose sections *Absus* and *Chamaecrista*, including those that had their bacteroids enclosed in either symbiosomes (Fig. 4E, S2J) or were intermediate (FT-SYM), such as *C. arrojadoana* and *C. mucronata* (Fig. 4F, S2L). It should be noted, however, that this was not a uniform observation across these *Chamaecrista* sections, as nodules of many species had no JIM5 labelling, such as *C. chapadae* (Fig. S2J), and of those that did have it the JIM5 epitope was located within the symbiosomes themselves (Fig. 4E). Image analysis of the infected N-fixing cells in sections from nodules on the species shown in Fig. S2 indicated that the FT-type nodules from section *Apoucouita* were less densely colonized by bacteroids than those of the SYM-type nodules in sections *Absus* and *Chamaecrista*, while the FT-SYM-type nodules from section *Baseophyllum* were intermediately colonized (Fig. S3).

Light microscopy combined with immunohistochemistry using polyclonal antibodies against *P. phymatum* STM815^T^ and *Cupriavidus taiwanensis* LMG19424^T^ indicated the presence of *Paraburkholderia* as symbionts in nodules of the tree species *C. eitenorum* (Fig. 5A, B). Specific immunogold localization of the *P. phymatum* antibody to the bacteroids was confirmed under the TEM (Fig. 5C, D). Further immunogold localization with JIM5 revealed HG epitopes on the cell walls surrounding both the FTs (Fig. 5E) and the invasive ITs (5F) in the *C. eitenorum* nodules containing their *Paraburkholderia* symbionts.

**Fig. 5.**
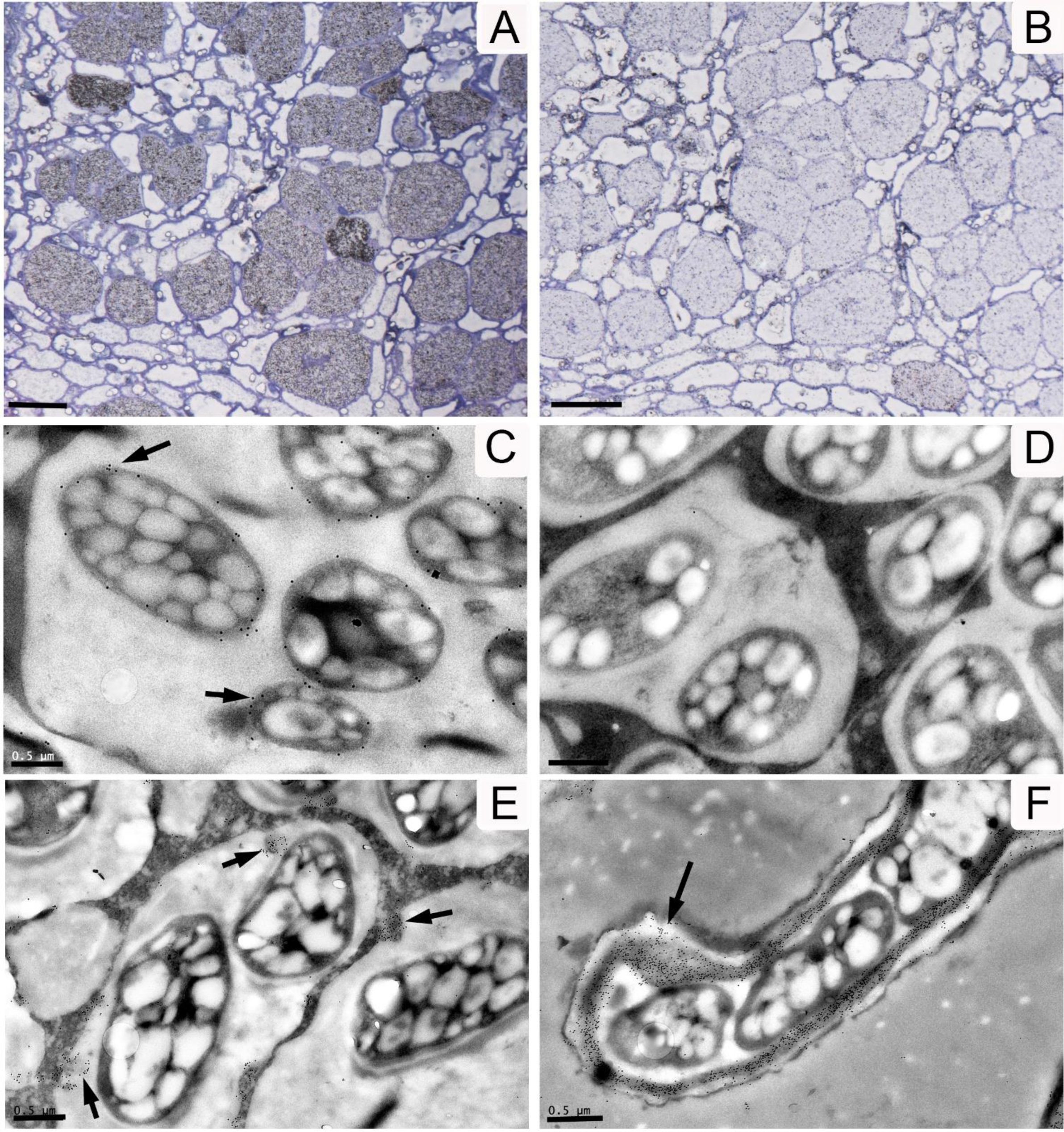
Light micrographs (**A, B)** and TEMs (**C – F**) of sections of *Chamaecrista eitenorum* var. *eitenorum* (Section *Apoucouita*) nodules. **A.** Infected tissue immunogold labelled with an antibody against *Paraburkholderia phymatum* STM815^T^ plus silver enhacement; **B.** Infected tissue immunolabelled with an antibody against *Cupriavidus taiwanensis* LMG19434^T^ plus silver enhancement (negative control). **C.** Immunogold localization on the bacteroid surface using an antibody against *Paraburkholderia phymatum* STM815^T^ (arrows). **D.** Negative control using IGL buffer. **E.** Immunogold localization of homogalacturonan epitopes (arrows) with the monoclonal antibody JIM5, which recognizes a pectin epitope in plant cell walls, in an infected cell. **F.** A “classical” invasion infection thread strongly immunogold labelled with JIM5 (arrows). Scale bars: **A, B** = 50 μm, **C, D, E, F** = 0.5 μm.

### Phylogenetic analysis of nitrogen-fixing symbionts: housekeeping genes and ITS region

The diversity of rhizobia nodulating the various *Chamaecrista* species in the present study was examined by MLSA to establish the phylogenetic relationships between symbiont strains associated with tree and treelet *Chamaecrista* species *vis-a-vis* strains deposited in the databases plus those isolated in our earlier study of symbionts from subshrub *Chamaecrista* spp. (Santos et al. 2017) (Fig. 6). Most strains were grouped in the genus *Bradyrhizobium* on the basis of close similarity to sequences of type strains. This genus comprises various supergroups, including two mega-clades: I, the *B. japonicum* group, and II, the *B. elkanii* group (Menna et al. 2009; Avontuur et al. 2019; Ormeno-Orillo and Martinez-Romero, 2019). One cluster of *Chamaecrista* strains (Tree cluster I; Fig. 6) consisting of isolates from the trees *C. ensiformis* var. *plurifoliolata, C. bahiae* and *C. duartei* were located within the *B. japonicum* supergoup, while the other cluster of *Chamaecrista* tree strains (Tree cluster II; Fig. 6), consisting only of isolates from *C. ensiformis* var. *plurifoliolata* were located within the *B. elkanii* supergroup. Mega-clade I also contained a large cluster of strains formed exclusively of bradyrhizobia isolated by Santos et al. (2017) from root nodules of subshrub *Chamaecrista* species (Subshrub-Shrub cluster; Fig. 6).

**Fig. 6.**
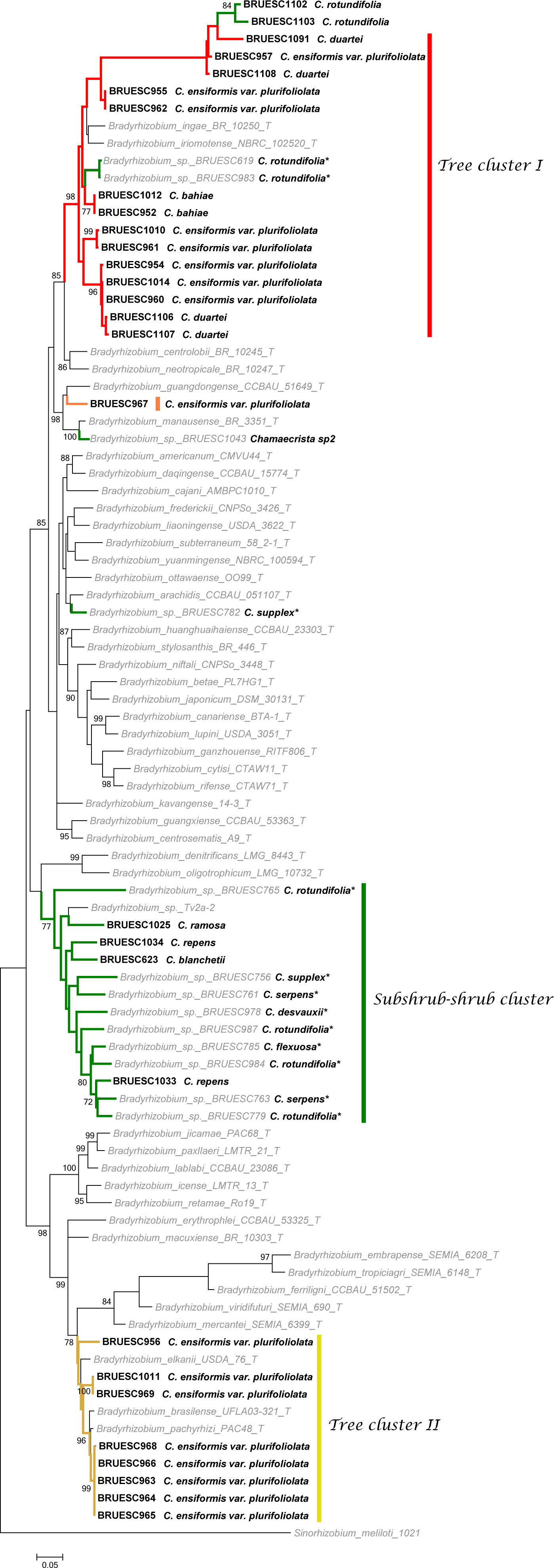
Maximum-likelihood phylogeny for the genus *Bradyrhizobium* based on the concatenated dataset consisting of sequences of the genes *atpD, dnaK, glnII, gyrB, recA* and *rpoB*. The isolates examined in this study are indicated in bold, and those from dos Santos et al. (2017) are indicated by *. The scale bar indicates the number of nucleotide substitutions per site. Numbers on branches are bootstrap values for 1000 replications (shown only when ≥70%).

Analysis of the ITS region (16S–23S rDNA) (Fig. S4) suggested congruency with the MLSA phylogeny, with two clusters containing strains from nodules of tree *Chamaecrista* spp. plus one Subshrub-Shrub cluster. Strains from subshrubs formed a separate cluster from all the *Bradyrhizobium* groups (Fig. S4). The ITS analysis indicates the large differences in the DNA sequences of these *Chamaecrista* isolates, as it is one of the strongest tools for discriminating between bradyrhizobial populations owing to its high degree of variation providing greater powers of resolution (Willems et al., 2003). In general, the *rrs* (16S rRNA) phylogeny (Fig. S5) was congruent with the MLSA and ITS phylogenies, but with less precision and with modification in the position of some bradyrhizobial isolates. None of the Bahia *Chamaecrista* strains clustered with the described species from *C. fasciculata*, *B. frederickii* (Urquiaga et al., 2019) and *B. niftali* (Klepa et al., 2019), both of which were isolated in the USA.

Confirming the immunogold microscopy observations for this species (Fig. 5A – D), both strains isolated from nodules on the tree *C. eitenorum* sampled in Chapada Diamantina belonged to the genus *Paraburkholderia*. A concatenated (16S rDNA + *recA*) phylogenetic analysis demonstrated that they are potentially a new species (Fig. S6) related to *P. nodosa* that was originally isolated from nodules of *Mimosa scabrella* (Chen et al., 2007).

### Phylogenetic analysis of nitrogen-fixing symbionts: symbiotic genes

The *nodC* (Fig. 7) and *nifH* (Fig. S7) bradyrhizobia phylogenies were relatively congruent with the concatenated housekeeping and ITS phylogenies, with both forming the same two Tree I and Tree II clusters as the MLSA, but the single subshrub-shrub cluster (SSC) comprising strains from this study and that of Santos et al. (2017) was divided into two sub-groups (SSCI & SSCII) in the *nodC* phylogeny (Fig. 7). SSCI corresponded to Cluster 1 of Santos et al. (2017) but now also incorporated *C. rotundifolia* and *C. ramosa* symbionts from the present study, while SSCII was more heterogenous, grouping several Santos et al. (2017) strains with a *C. blanchetii* strain from the present study (BRUESC623) plus strains from various other legumes. A single strain (BRUESC1034) isolated from the shrub *C. repens* occupied a unique position outside any of the main *nodC* clusters. For the tree symbionts, Tree cluster II (TCII) constituted a tight group of six strains from *C. ensiformis* var. *plurifoliolata*, while Tree cluster I (TCI) constituted the main group of *Chamaecrista* tree strains; these were related to *Bradyrhizobium* type species that were isolated in Brazil from various legumes (except for *B. iriomotense* EK05^T^ from Japan). The *nifH* phylogeny differed from the *nodC* one in that the two SSC sub-groups were not apparent, with all the strains being grouped into a single cluster with *Bradyrhizobium* sp. Tv2a-2 2 (isolated from the caesalpinioid legume *Tachigali versicolor* in Barro Colorado Island of Panama; Tian et al., 2015) and *B. ganzhouense*, a symbiont of *Acacia melanoxylon* (Lu et al. 2014).

**Fig. 7.**
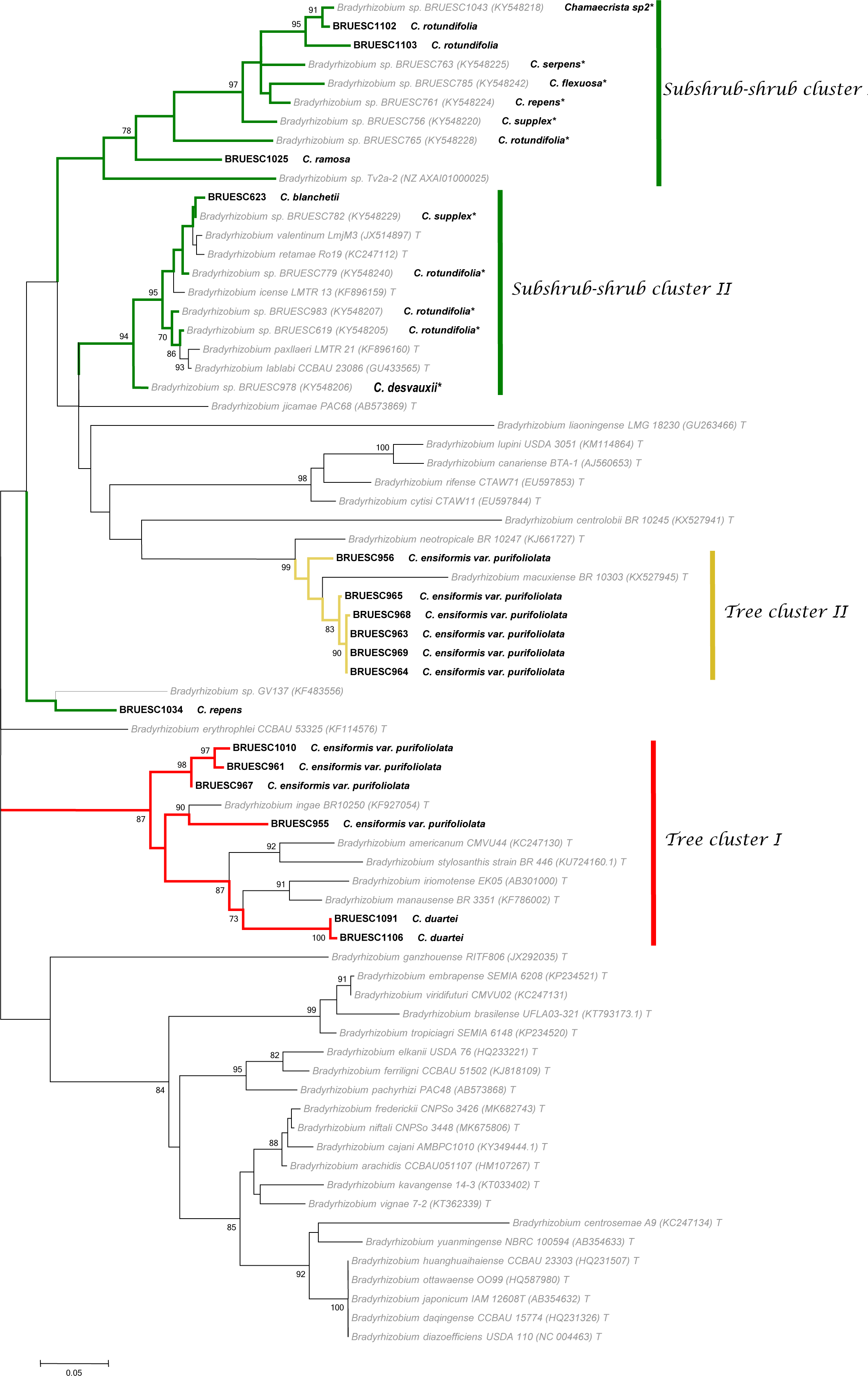
Maximum-likelihood phylogeny of the genus *Bradyrhizobium* based on *nodC*. Host association and specific geographic origin is listed in Table 1. The isolates examined in this study are indicated in bold, and those from dos Santos et al. (2017) are indicated by *. The scale bar indicates the number of nucleotide substitutions per site. Numbers on branches are bootstrap values for 1000 replications (shown only when ≥70%).

The two strains of *Paraburkholderia* isolated from *C. eitenorum* (BRUESC1092, BRUESC1093) formed a tight cluster in the *nodC* phylogeny with *Paraburkholderia* sp. BRUESC684 (Fig. S8), a strain isolated from *Calliandra viscidula* collected in Chapada Diamantina in Bahia State (Silva et al., 2018), but it should also be noted that both *C. eitenorum* strains had symbiosis gene sequences which were closely related to *P. nodosa* and *P. mimosarum* which are widely isolated symbionts of *Mimosa* species in Brazil (Bontemps et al., 2010). Strains BRUESC1092 and BRUESC1093 also grouped with the *P. nodosa* type strain Br3437^T^ in the *nifH* gene phylogeny (Fig. S9).

### Nodulation ability and host range of the Chamaecrista rhizobia

From the 28 strains of *Bradyrhizobium* isolated from nodules of seven *Chamaecrista* species (*C. bahiae*, *C. duartei*, *C. eitenorum, C. ensiformis* var. *plurifoliolata, C. blanchetii*, *C. ramosa*, *C. repens* and *C. rotundifolia*) 23 were tested for their nodulation ability on Siratro, and 15 were tested on six *Chamaecrista* spp. The tested strains represented the whole range of growth habits in the genus, from trees to subshrubs (Table 2), but also the four *nodC* genotype clusters identified in Fig. 7 (SSCI, SSCII, TCI and TCII). Five of the strains were isolated from the tree *C. ensiformis* var. *plurifoliolata*; three of these, representing *nodC* TCI (BRUESC1010) and TCII (BRUESC964), and BRUESC1011 (not included in the *nodC* phylogeny) formed nodules on their homologous host (the other two strains, BRUESC956 and BRUESC967 were not tested on *C. ensiformis*), and four of the five strains nodulated the subshrub *C. rotundifolia* (BRUESC964 was not tested). However, no nodules were observed when one of the *C. ensiformis* var. *plurifoliolata* strains (BRUESC1011) was inoculated onto another *Chamaecrista* tree species (*C. bahiae*). A *Bradyrhizobium* strain isolated from *C. bahiae* (BRUESC952) nodulated *C. rotundifolia,* but not *C. blanchetii* and *C. desvauxii*, while strain BRUESC1107 from *C. duartei* formed effective nodules on *C. desvauxii*. The majority of the 19 *Bradyrhizobium* strains tested nodulated Siratro; the exceptions were BRUESC623 from *C. blanchetii* BRUESC964 from *C. ensiformis* var. plurifoliolata, BRUESC1033 and BRUESC1034 from *C. repens*, and BRUESC1106 from *C. duartei* (Table 2). Finally, the two *Paraburkholderia* strains isolated from the tree *C. eitenorum* (BRUESC1092 and BRUESC1093) effectively nodulated other tree species in the section *Apoucouita*, *C. duartei* and *C. ensiformis*, but nodulated *Mimosa pudica* ineffectively, and both failed to nodulate Siratro.

### Phylogenetic analysis of the genus Chamaecrista

The individual phylogenetic analyses of *Chamaecrista* did not show apparent incongruence between the ITS and *trnL-F* regions (Fig. 8), but as already described for *Chamaecrista* (Conceição et al., 2009; Rando et al., 2016), the ITS region gives higher resolution within the sections. In the *Chamaecrista* phylogeny (Fig. 8), three main large clades emerged with high support (PP=1). These groups corresponded to the section *Apoucouita* embracing the arborescent rainforest species, the section *Baseophyllum* consisting of treelets and shrubs from Campo Rupestre and Caatinga, and the sections *Absus* and *Chamaecrista*, which contain the highest diversity of species in the genus, with most species being shrubs and subshrubs. Interestingly, these main clades highlighted here are congruent with the phylogeny of the nitrogen-fixing microsymbionts, and also demonstrate that the FT phenotype in which bacteroids are enclosed in cell wall material, is apparently confined to the arborescent species in section *Apoucouita*, whereas in species from the other clades/sections the microsymbionts are mainly contained within symbiosomes. Interestingly, however, there are also a number of species, particularly in section *Baseophyllum*, but also scattered amongst the other sections, that are intermediate (FT-SYM) with regard to their bacteroid phenotype (Fig. 8).

**Fig. 8.**
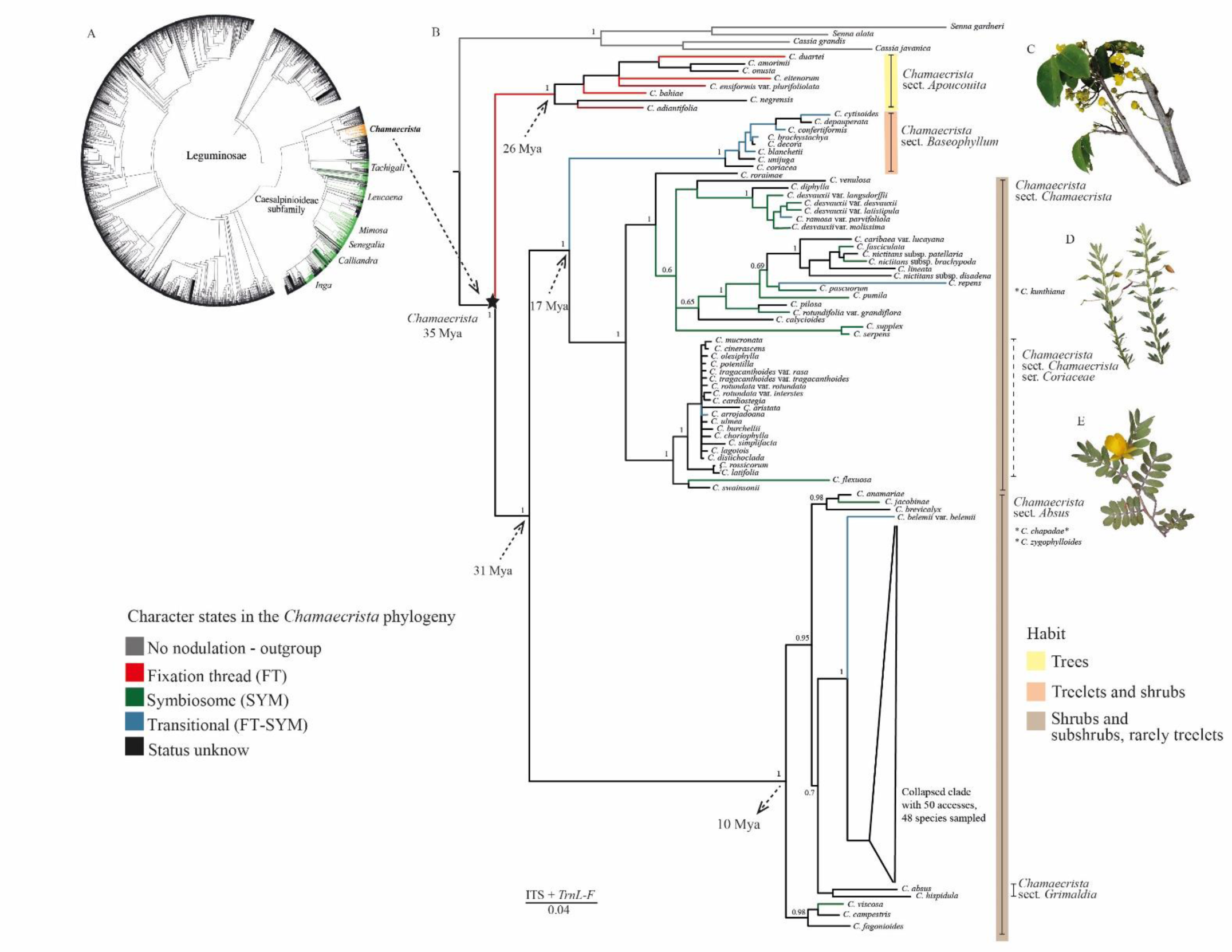
**A.** Phylogeny of the Leguminosae family adapted from LPWG (2017) showing the genera in the Caesalpinioideae subfamily associated with symbionts. **B.** Phylogeny of *Chamaecrista* based on DNA sequences of nuclear ITS and plastidial *trnL-trnF* loci. Majority-rule consensus tree derived from Bayesian analysis; the values of posterior probability (PP) are indicated above the branches in decimal form. Taxonomic groups of associated symbionts are colored. Species indicated by *(*C. chapadae*, *C. zygophylloides*) were not included in the phylogenetic analysis, but based on morphology, it was possible to place them in their appropriate taxonomic group. Arrows indicate the ages of nodes as estimated by Rando et al. (2016). **C.** *C. bahiae*; **D.** *C. desvauxii*; **E.** *C. arrojadoana.* (photos Juliana Rando)

## Discussion

### *The occurrence of fixation threads (FTs), symbiosomes or intermediates depends on the taxonomy, growth habit, and habitat of the* Chamaecrista *host*

The 26 new reports in the present study raises the number of confirmed nodulating *Chamaecrista* species to 74 (Sprent, 2009; Santos et al. 2017; Faria et al. 2022; Table S4), now representing nearly a quarter of this highly speciose genus. In general, tissue distribution in the interior of *Chamaecrista* nodules is similar to that of indeterminate nodules from other neotropical Caesalpinioid trees, such as *Anadenanthera peregrina* (Gross et al., 2002), *Mimosa* spp. (dos Reis Junior et al., 2010), and *Dimorphandra* spp. (Fonseca et al., 2012). However, the present study has also confirmed that *Chamaecrista* nodules are unique within the Caesalpinioideae since their infected cells can, depending on species, have their bacteroids either retained within FTs, as occurs in nodules on all other non-Mimosoid nodulated Caesalpinioideae, or have their rhizobial symbionts released into symbiosomes (SYM-type nodules), as occurs in all mimosoid and most papilionoid nodules so far reported (Naisbitt et al., 1992; Sprent, 2001; Sprent 2009; Sprent et al., 2017; Faria et al. 2022). The distribution of FT-versus SYM-type nodules in *Chamaecrista* appears to depend on the growth habit of the plant *i.e.,* current knowledge, reinforced by considerable additional data from the present study, indicates that tree species are all FT-type, whereas smaller shrub, subshrub and herbaceous species, which represent the majority of *Chamaecrista* species, tend to be SYM-types (Naisbitt et al. 1992). Interestingly, the present study has also confirmed using SEM and TEM that some treelet and large shrub species exhibit an intermediate (FT-SYM) type of nodule in which the FTs are less distinct (Naisbitt et al. 1992), suggesting a transitional stage between FTs and symbiosomes in *Chamaecrista* that has not so far been observed in other non-Mimosoid grade Caesalpinioideae symbioses (Faria et al. 2022).

### Composition of the FT in Chamaecrista species

As with other non-mimosoid Caesalpinioideae trees, such as *Dimorphandra* (Fonseca et al., 2012), *Erythrophleum* and *Moldenhawera* species (De Faria et al. 2022), the FTs in *Chamaecrista* tree species comprise a cell wall containing homogalacturonan (HG) epitopes as recognized by the monoclonal antibody, JIM5, which labels unesterified pectin. The present study has demonstrated that *Chamaecrista* FTs also contain cellulose; this concurs with the study of Faria et al. (2022) who have recently demonstrated that the cell wall of FTs in nodules on a range of non-mimosoid Caesalpinioideae contain several pectic components, but also cellulosic ones. Indeed, there is mounting evidence that the wall of the FT differs little from the IT cell wall (and plant cell walls in general) except that the FT wall is generally thinner and contains less unesterified pectin (JIM5 epitope) (Fonseca et al. 2012; Faria et al. 2022; this study). It should also be noted that FTs are not simple cell wall-bound structures, but are similar to “conventional” symbiosomes in possessing a host membrane that surrounds the cell wall (Faria et al. 2022), and hence the relative absence of unesterified pectin in FTs, as well as their relatively thin walls compared to ITs is likely to be an adaptation to allow the easier exchange of nutrients and O_2_ between the rhizobial bacteroids and the host cytoplasm across this membrane (Fonseca et al. 2012; Faria et al. 2022). Such nutrient and gaseous exchange would most likely be impeded by the thick cell wall of a typical IT containing pectin that has been stiffened via cross-linking through the action of pectin methylesterases (PME) (Su, 2023).

Our study has also revealed more about the ultrastructure of the FT-SYM- and SYM-type *Chamaecrista* nodules, including the histochemical nature of the symbiosomes. For example, JIM5-labelled material could be observed in a matrix that surrounds the symbiosomes in mature infected cells from nodules of some SYM and FT-SYM-type *Chamaecrista* shrub and subshrub species in the present study from Brazil, but also in the Asian subshrub species *C. pumila* (Rathi et al. 2018). The JIM5 labelling observed in the FT-SYM-type nodules is most likely because these are essentially thinner versions of FTs, and still have obvious cell walls surrounding their bacteroids (Naisbitt et al. 1992; this study), but it is curious that some SYM-type nodules also possess pectin without any obvious walls. In pea nodules, although glycoproteins and glycolipids were observed, symbiosomes contained neither polysaccharides nor cell wall material (Perotto et al., 1991, 1995). This implies that our observations of bacteroids in some SYM-type *Chamaecrista* nodules surrounded by partially methyl-esterified and unesterified HG epitopes (JIM5) are different from the symbiosomes housing bacteroids within nodules on species in the “advanced” Inverse Repeat-Lacking Clade (IRLC); the latter contain greatly-enlarged endo-reduplicated bacteroids that have completely lost their ability to divide and to proliferate outside the host plant i.e. they are essentially organelles (Brewin, 1991; Gage, 2004; Oono et al. 2010; Sprent et al. 2017; Ardley & Sprent, 2021; Mathesius, 2022). On the other hand, such terminally differentiated bacteroids are not the norm outside the IRLC, so they are clearly not essential for legume nodule functioning, although they may be more efficient at fixing N_2_ (Mathesius, 2022). In most other legume nodules, undifferentiated bacteroids are housed within symbiosomes, and it could be argued that the *Chamaecrista* SYM-type nodules are essentially like these, especially those in the Mimosoid clade (De Faria et al. 2022 and references therein). Therefore, as suggested in the earlier study on a much narrower range of species (Naisbitt et al. 1992), it would indeed appear that *Chamaecrista* contains a range of nodule phenotypes from full FTs in the trees in the section *Apoucouita* through intermediates (FT-SYM) in treelets and large shrubs (e.g. in the section *Baseophyllum*, although not exclusively) to “standard” symbiosomes (SYM-type) in the majority of *Chamaecrista* species in the sections *Absus* and *Chamaecrista*. The comparative efficiencies of the different nodule types to fix N are not yet clear, but the number of bacteroids per cell (Fig. S3, this study) are significantly reduced in the FT-type compared to the SYM-type nodule, with the FT-SYM-types being intermediate in both parameters, which suggests that the possession of a cell wall around the bacteroids reduces the degree to which host cells can be packed with N-fixing rhizobia. This, together with and reduced proportion of infected cells per nodule (Naisbitt et al. 1992) is likely to reduce the overall efficiency of the FT-type nodule.

### The influence of plant habit and biome on the diversity of nitrogen-fixing symbionts in Chamaecrista

Our study plus that of Santos et al. (2017) has demonstrated that in common with almost all non-Mimosoid legumes of the subfamily Caesalpinioideae (Fonseca et al., 2012; Parker, 2015; Cabral Michel et al., 2021; Avontuur et al. 2021), symbiotic bacteria isolated from nodules across the genus *Chamaecrista* belong almost exclusively to the genus *Bradyrhizobium*. Both housekeeping genes and ITS sequences of *Chamaecrista* strains were congruent and taken together with the two symbiosis related genes (*nifH* and *nodC*) showed that at least a dozen strains could represent putative novel species of *Bradyrhizobium*. Also, *C. ensiformis,* a widespread arborescent species from tropical forests, had the highest diversity of *Bradyrhizobium* symbionts, differing from its closely-related “cousin” in the section *Apoucouita*, *C. eitenorum,* which has a more restricted distribution in SDTF.

The phylogenetic analysis of both the *nodC* and *nifH* genes showed that the *Bradyrhizobium* strains obtained from *Chamaecrista* nodules sampled in Bahia formed clearly separated branches suggesting that symbiotic genes were probably vertically transmitted, as was already indicated by studies on bradyrhizobia from other legumes (Moulin et al., 2004; Parker, 2015, Fonseca et al., 2012; Stepkowski et al., 2007, 2018). These results also suggest a very high diversity of bradyrhizobia nodulating *Chamaecrista* trees, shrubs and subshrubs native to Brazil, and to Bahia in particular. The *nodC* phylogeny supported the likelihood of there being co-evolution and specificity of the *Chamaecrista-Bradyrhizobium* symbiosis since the rhizobia of the subshrub *Chamaecrista* species in sections *Absus* and *Chamaecrista* that are highly divergent from their distant cousins in the rainforest tree section *Apoucouita* also had highly divergent *nodC* sequences. On the other hand, the different environments that these plant species occupy, especially their soils (e.g. pH), will also play a part in their selection of particular symbionts, as demonstrated for *Mimosa* in the Cerrado (Pires et al. 2018), and for *Chamaecrista* in India (Rathi et al. 2018). Indeed, although the cross-inoculation experiments demonstrated some degree of specificity, particularly for *Chamaecrista* tree and treelet species, the fact that the widely distributed subshrub *C. rotundifolia* is nodulated by *Bradyrhizobium* strains from all the observed *nodC* clades, including those from the section *Apoucouita* species *C. bahiae* and *C. ensiformis*, as well as the ability of most of the tested isolates to nodulate Siratro, suggests that Brazilian *Chamaecrista* symbionts can be cosmopolitan in their selection of hosts outside their normal native ranges, despite their apparent co-evolution with their (often) endemic hosts. A similar observation was made with *Mimosa*, another large nodulated legume genus that has radiated in Bahia, albeit with Beta-rhizobial symbionts rather than bradyrhizobia (Bontemps et al. 2010).

Although bradyrhizobia are clearly the principal symbionts of *Chamaecrista* in Brazil and elsewhere (see Introduction), other rhizobial types can nodulate the genus, such as *Sinorhizobium* (*Ensifer*) in alkaline soils in India (Rathi et al. 2018). In the present study, the tree species *C. eitenorum*, which is native to the SDTF, had the apparently unique property of being nodulated by strains of *Paraburkholderia* related to those that nodulate *Mimosa* and *Calliandra* spp. in the same environment (Bontemps et al. 2010; Silva et al. 2018); indeed, the fact that these isolates were also capable of nodulating *C. duartei* and *C. ensiformis* in cross-inoculation studies suggests that *Chamaecrista*-*Paraburkholderia* symbioses might be relatively abundant, at least among the tree species. Further to this, the *C. eitenorum-Paraburkholderia* interaction is the first confirmed report of a Beta-rhizobial symbiosis in nodules from a Caesalpinioideae species outside the Mimosoid clade (LPWG, 2017), and it is also the first report of FTs being occupied by this type of microsymbiont, which demonstrates that the FT phenotype is not controlled by any particular rhizobial type (e.g. *Bradyrhizobium*), but is an inherent plant characteristic, which might be expected if the “host controls the party” (Ferguson et al. 2019).

### Concluding remarks

Our molecular phylogeny of *Chamaecrista* exhibits the same relationship among the monophyletic clades already observed in other studies (Conceição et al., 2009; Rando et al., 2016; Mendes et al., 2020; Souza et al., 2021), even with the incorporation of new taxa. The small section *Apoucouita* remains a monophyletic group with the addition of six new arborescent species. The section also appears as a sister group of all remaining species of the genus, corroborating that its habitat and habit diverged distinctly from the others, and that it may be basal within the genus. The shifts of growth habit appear to be correlated with the habitat changes in *Chamaecrista*. In the genus, the diversification occurred from rain forests with arborescent habits to open areas (savannahs and SDTF) with smaller shrub and subshrub growth habits (Conceição et al., 2009; Coutinho et al. 2016; Souza et al., 2021).

We propose that the lower nutrient soils (and several other stresses, such as seasonal drought) associated with savannahs and SDTF not only drove a reduction in growth habit, but also necessitated a greater dependency on a more reliable nodulating symbiosis, as the smaller root systems of shrub and subshrub *Chamaecrista* species have a reduced access to a much more limited pool of soil N compared to large rainforest-dwelling trees that also have a higher capacity to (re)cycle their N. Therefore, the shrub and subshrub *Chamaecrista* species have largely rejected the FT-type symbiosis of their arboreal cousins in the section *Apoucouita*, and have gradually adopted (via the intermediate FT-SYM-type nodule) the more intimate SYM-type nodule, which with its full incorporation of the bacteroids into the host cytoplasm exhibits the compartmentalization that is linked with more stable and efficient symbioses (Parniske, 2018; Chomicki et al. 2020; Faria et al. 2022; Libourel et al. 2023; James 2023; Mohd-Radzman & Drapek, 2023). In this respect, the FT versus SYM pattern revealed by the present study across the highly speciose genus *Chamaecrista* mirrors that across the Caesalpinioideae subfamily as a whole i.e. that the hugely diverse (in terms of both growth habits and number of genera) Mimosoid clade has retained nodulation by rejecting the less intimate and relatively unstable FT-type nodule of its few nodulating cousins in the Caesalpinioideae grade that subtends it (with the notable exception of *Chamaecrista*), and by adopting the SYM-type nodule has avoided the massive losses of nodulation evident in the non-Mimosoid Caesalpinioideae (Faria et al. 2022).

## Supporting information

Table S2 & S4 Casaes et al.

Suppl Material Casaes et al.

## ACKNOWLEDGEMENTS

The authors thank Dr Alan Prescott and Tamanatha Jithesh for assistance with image analysis, and Professor A. Nogueira and Dr Chrizelle Beukes for critical reading and suggestions. We would like to thank FAPESB for a grant to PAC Alves via the postgraduate Program of Biology and Biotechnology of Microorganisms, and the Center for Electron Microscopy at UESC for infrastructure. EK James acknowledges funding from CAPES via the Ciençia sem Fronteiras programme.

---

**Fig. S1.** Location and distribution of *Chamaecrista* nodule sampling in Bahia State (NE Brazil).

**Fig. S2.** Light micrographs (**A, C, E, G, I, K**) and transmission electron micrographs (TEMs) plus immunogold labelling with JIM5 (**B, D, F, H, J, L**) of nodules of *Chamaecrista* across four sections of the genus illustrating the range of nodule anatomical types from tree species with their bacteroids enclosed in fixation threads (FT) labelled with JIM5 (**A – D**) through intermediate types (FT-SYM) on treelets (**E – H**) to membrane-bound symbiosomes (SYM) on shrub and subshrub species that have little or no JIM5 signal (**I – L**). **A, B.** *Chamaecrista duartei* (section *Apoucouita*). **C, D.** *Chamaecrista ensiformis* (section *Apoucouita*). **E, F.** *Chamaecrista blanchettii* (section *Baseophyllum*). **G, H.** *Chamaecrista cytisoides* (section *Baseophyllum*). **I, J.** *Chamaecrista chapadae* (section *Absus*, subsect. *Absus*). **K, L.** *Chamaecrista mucronata* (section *Chamaecrista*). The FTs in the section *Apoucouita* tree species *C. duartei* (**A**) and *C. ensiformis* (**C**) are discernible in infected cells (*) at the light microscope level. The walls of the FTs are indicated by arrows in TEMs in **B** and **D,** but note that the FTs in *C. ensiformis* (**D**) are more densely labelled with JIM5 than those in *C. duartei* (**B**). However, in the latter species the electron dense cell walls are more apparent, thus illustrating that there is no direct relationship between cell wall density and the presence/absence of unesterified pectin (JIM5) in FTs. Bacteroids in nodules on the section *Baseophyllum* treelet species *C. blanchetii* (**E**) and *C. cytisoides* (**G**) are less clearly defined at the light microscope level and this is because at the TEM level (**F, H**) it is observed that bacteroids either have thin FT walls with sparse JIM5 labelling (arrows in **F, H**) or no walls at all (i.e., most of the bacteroids in *C. cytisoides*, **H**); however, more densely labelled walls surrounding bacteroids were also occasionally observed within *C. cytisoides* infected cells (double arrows in **H**), suggesting that this species harbours both FTs and symbiosomes. The bacteroids in nodules on shrub and subshrub species *C. chapadae* (**I**) and *C. mucronata* (**K**) were even less distinct under the light microscope compared to the *Baseophyllum* treelet species, and this was also confirmed under the TEM where they were shown to be enclosed in symbiosomes Bars = 20 µm (**A, C, E, G. I, K**), 500 nm (**B, D, F, L**), 1 µm (**H, J**), b = bacteroid, w = plant cell wall in **B, D, F, H, J, L.**

**Fig. S3.** Proportional (%) occupation of infected cells by symbiotic rhizobia estimated by counting pixels from light micrographs similar to those presented in Fig S2A and C for FT-type, Fig. S2E and G for FT-SYM-type, and Fig. S2I and K for SYM-type nodules. Data are presented as the mean proportion (± s.d.) of each infected cell profile filled with toluidine blue-stained structures representing bacteroids; 7-9 sections were examined per symbiosome type (FT, FT-SYM, SYM).

**Fig. S4.** Maximum-likelihood phylogeny for the genus *Bradyrhizobium* based on the ITS region. Host associations and specific geographic origins are listed in Table 1. The isolates examined in this study are indicated in bold, and those from dos Santos et al. (2017) are indicated by *. The scale bar indicates the number of nucleotide substitutions per site. Numbers on branches are bootstrap values for 1000 replications (shown only when ≥70%).

**Fig. S5.** Maximum-likelihood phylogeny for the genus *Bradyrhizobium* based on 16S rDNA. Host association and specific geographic origin is listed in Table 1. The isolates examined in this study are indicated in bold, and those from dos Santos et al. (2017) are indicated by *. The scale bar indicates the number of nucleotide substitutions per site. Numbers on branches are bootstrap values for 1000 replications (shown only when ≥70%).

**Fig. S6.** Maximum-likelihood phylogeny for the genus *Paraburkholderia* based on the concatenated dataset consisting of sequences for the genes 16S rDNA and *recA*. The isolates examined in this study are indicated in bold. The scale bar indicates the number of nucleotide substitutions per site. Numbers on branches are bootstrap values for 1000 replications (shown only when ≥70%).

**Fig. S7.** Maximum-likelihood phylogeny for the genus *Bradyrhizobium* based on the *nifH* gene. Host association and specific geographic origin is listed in Table 1. The isolates examined in this study are indicated in bold, and those from dos Santos et al. (2017) are indicated by *. The scale bar indicates the number of nucleotide substitutions per site. Numbers on branches are bootstrap values for 1000 replications (shown only when ≥70%).

**Fig. S8.** Maximum-likelihood phylogeny for the genus *Parburkholderia* based on the *nodC* gene. Host association and specific geographic origin is listed in Table 1. The isolates examined in this study are indicated in bold. The scale bar indicates the number of nucleotide substitutions per site. Numbers on branches are bootstrap values for 1000 replications (shown only when ≥70%).

**Fig. S9.** Maximum-likelihood phylogeny for the genus *Paraburkholderia* based on the *nifH* gene. Host association and specific geographic origin is listed in Table 1. The isolates examined in this study are indicated in bold. The scale bar indicates the number of nucleotide substitutions per site. Numbers on branches are bootstrap values for 1000 replications (shown only when ≥70%).

**Table S1.** Accession numbers of aerial parts of *Chamaecrista* species sampled in Bahia (BA), Brazil and deposited in the Herbarium of the Universidade Estadual de Santa Cruz (UESC). Also included is information about growth habit, vegetation type and locality where the *Chamaecrista* species were sampled in BA specifically for this study.

**Table S2.** Genbank accession numbers of gene sequences from strains isolated in the present study.

**Table S3.** Voucher information and GenBank accession numbers of the nuclear ITS and plastidial *trnL-trnF* loci sequences included in the phylogeny of *Chamaecrista* (Fig. 8). New sequences are marked with an asterisk (*).

**Table S4.** Confirmed positive global nodulation reports for *Chamaecrista*; data are taken from Sprent (2009) unless noted otherwise. Growth habit and habitat information were either noted by the authors during sample collection in Brazil or were extracted from *Flora e Funga do Brasil* (https://floradobrasil.jbrj.gov.br/FB22876) and/or *Plants of the World Online* (https://powo.science.kew.org/). Nodule anatomy data, including the presence of either Symbiosome (SYM), Fixation Threads (FT), or Intermediate (FT-SYM) are taken from the literature, from the study of Sprent et al. (1996), or from the present study. Taxonomic information with regards to Sections and Subsections were extracted from Souza et al. (2021).

## Notes

### Competing Interest Statement

The authors have declared no competing interest.

### Summary of Updates

Legends added below figures

