## Supplementary material for "The radiation of nodulated *Chamaecrista* species from the rainforest into more diverse habitats has been accompanied by a reduction in growth form and a shift from fixation threads to symbiosomes": Suppl Material Casaes et al.

**Fig. S1.** Location and distribution of *Chamaecrista* nodule sampling in Bahia State (NE Brazil).

**Fig. S2.** Light micrographs (**A, C, E, G, I, K**) and transmission electron micrographs (TEMs) plus immunogold labelling with JIM5 (**B, D, F, H, J, L**) of nodules of *Chamaecrista* across four sections of the genus illustrating the range of nodule anatomical types from tree species with their bacteroids enclosed in fixation threads (FT) labelled with JIM5 (**A – D**) through intermediate types (FT-SYM) on treelets (**E – H**) to membrane-bound symbiosomes (SYM) on shrub and subshrub species that have little or no JIM5 signal (**I – L**). **A, B.** *Chamaecrista duartei* (section *Apoucouita*). **C, D.** *Chamaecrista ensiformis* (section *Apoucouita*). **E, F.** *Chamaecrista blanchettii* (section *Baseophyllum*). **G, H.** *Chamaecrista cytisoides* (section *Baseophyllum*). **I, J.** *Chamaecrista chapadae* (section *Absus*, subsect. *Absus*). **K, L.** *Chamaecrista mucronata* (section *Chamaecrista*). The FTs in the section *Apoucouita* tree species *C. duartei* (**A**) and *C. ensiformis* (**C**) are discernible in infected cells (*) at the light microscope level. The walls of the FTs are indicated by arrows in TEMs in **B** and **D,** but note that the FTs in *C. ensiformis* (**D**) are more densely labelled with JIM5 than those in *C. duartei* (**B**). However, in the latter species the electron dense cell walls are more apparent, thus illustrating that there is no direct relationship between cell wall density and the presence/absence of unesterified pectin (JIM5) in FTs. Bacteroids in nodules on the section *Baseophyllum* treelet species *C. blanchetii* (**E**) and *C. cytisoides* (**G**) are less clearly defined at the light microscope level and this is because at the TEM level (**F, H**) it is observed that bacteroids either have thin FT walls with sparse JIM5 labelling (arrows in **F, H**) or no walls at all (i.e., most of the bacteroids in *C. cytisoides*, **H**); however, more densely labelled walls surrounding bacteroids were also occasionally observed within *C. cytisoides* infected cells (double arrows in **H**), suggesting that this species harbours both FTs and symbiosomes. The bacteroids in nodules on shrub and subshrub species *C. chapadae* (**I**) and *C. mucronata* (**K**) were even less distinct under the light microscope compared to the *Baseophyllum* treelet species, and this was also confirmed under the TEM where they were shown to be enclosed in symbiosomes  Bars = 20 µm (**A, C, E, G. I, K**), 500 nm (**B, D, F, L**), 1 µm (**H, J**), b = bacteroid, w = plant cell wall in **B, D, F, H, J, L.**

**Fig. S3.** Proportional (%) occupation of infected cells by symbiotic rhizobia estimated by counting pixels from light micrographs similar to those presented in Fig S2A and C for FT-type, Fig. S2E and G for FT-SYM-type, and Fig. S2I and K for SYM-type nodules. Data are presented as the mean proportion (± s.d.) of each infected cell profile filled with toluidine blue-stained structures representing bacteroids; 7-9 sections were examined per symbiosome type (FT, FT-SYM, SYM).

**Fig. S4.** Maximum-likelihood phylogeny for the genus *Bradyrhizobium* based on the ITS region. Host associations and specific geographic origins are listed in Table 1. The isolates examined in this study are indicated in bold, and those from dos Santos et al. (2017) are indicated by *. The scale bar indicates the number of nucleotide substitutions per site. Numbers on branches are bootstrap values for 1000 replications (shown only when ≥70%).

**Fig. S5.** Maximum-likelihood phylogeny for the genus *Bradyrhizobium* based on 16S rDNA. Host association and specific geographic origin is listed in Table 1. The isolates examined in this study are indicated in bold, and those from dos Santos et al. (2017) are indicated by *. The scale bar indicates the number of nucleotide substitutions per site. Numbers on branches are bootstrap values for 1000 replications (shown only when ≥70%).

**Fig. S6.** Maximum-likelihood phylogeny for the genus *Paraburkholderia* based on the concatenated dataset consisting of sequences for the genes 16S rDNA and *recA*. The isolates examined in this study are indicated in bold. The scale bar indicates the number of nucleotide substitutions per site. Numbers on branches are bootstrap values for 1000 replications (shown only when ≥70%).

**Fig. S7.** Maximum-likelihood phylogeny for the genus *Bradyrhizobium* based on the *nifH* gene. Host association and specific geographic origin is listed in Table 1. The isolates examined in this study are indicated in bold, and those from dos Santos et al. (2017) are indicated by *. The scale bar indicates the number of nucleotide substitutions per site. Numbers on branches are bootstrap values for 1000 replications (shown only when ≥70%).

**Fig. S8.** Maximum-likelihood phylogeny for the genus *Parburkholderia* based on the *nodC* gene. Host association and specific geographic origin is listed in Table 1. The isolates examined in this study are indicated in bold. The scale bar indicates the number of nucleotide substitutions per site. Numbers on branches are bootstrap values for 1000 replications (shown only when ≥70%).

**Fig. S9.** Maximum-likelihood phylogeny for the genus *Paraburkholderia* based on the *nifH* gene. Host association and specific geographic origin is listed in Table 1. The isolates examined in this study are indicated in bold. The scale bar indicates the number of nucleotide substitutions per site. Numbers on branches are bootstrap values for 1000 replications (shown only when ≥70%).

**Table S1.** Accession numbers of aerial parts of *Chamaecrista* species sampled in Bahia (BA), Brazil and deposited in the Herbarium of the Universidade Estadual de Santa Cruz (UESC). Also included is information about growth habit, vegetation type and locality where the *Chamaecrista* species were sampled in BA specifically for this study.

**Table S2.** Genbank accession numbers of gene sequences from strains isolated in the present study. Excel file.

**Table S3.** Voucher information and GenBank accession numbers of the nuclear ITS and plastidial *trnL-trnF* loci sequences included in the phylogeny of *Chamaecrista* (Fig. 8). New sequences are marked with an asterisk (*).

**Table S4.** Confirmed positive global nodulation reports for *Chamaecrista*; data are taken from Sprent (2009) unless noted otherwise. Growth habit and habitat information were either noted by the authors during sample collection in Brazil or were extracted from *Flora e Funga do Brasil* (https://floradobrasil.jbrj.gov.br/FB22876) and/or *Plants of the World Online* (https://powo.science.kew.org/). Nodule anatomy data, including the presence of either Symbiosome (SYM), Fixation Threads (FT), or Intermediate (FT-SYM) are taken from the literature, from the study of Sprent et al. (1996), or from the present study. Taxonomic information with regards to Sections and Subsections were extracted from Souza et al. (2021). Excel file.

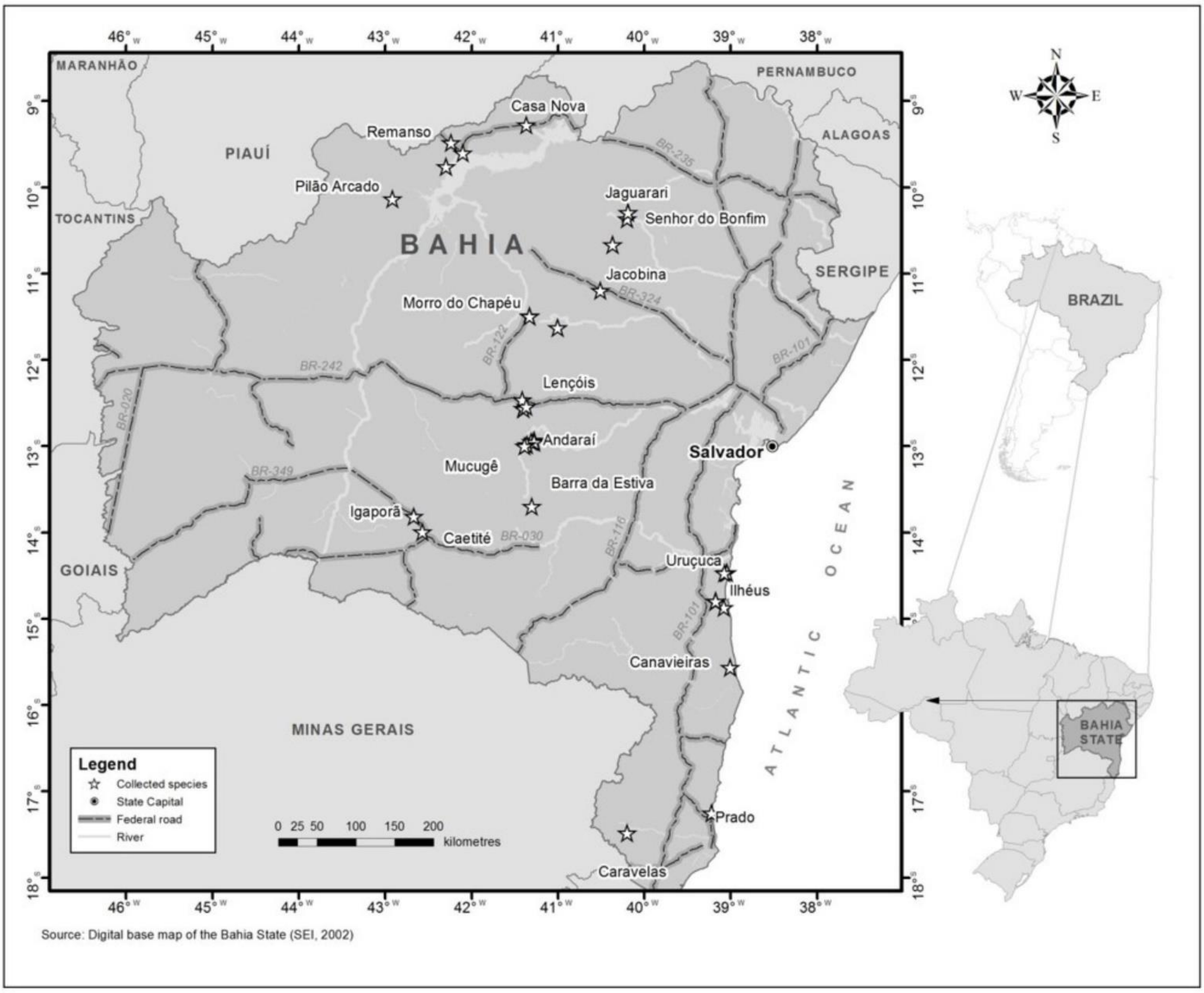


**FIG S1** Location and distribution of *Chamaecrista* nodule sampling in Bahia State (NE Brazil).

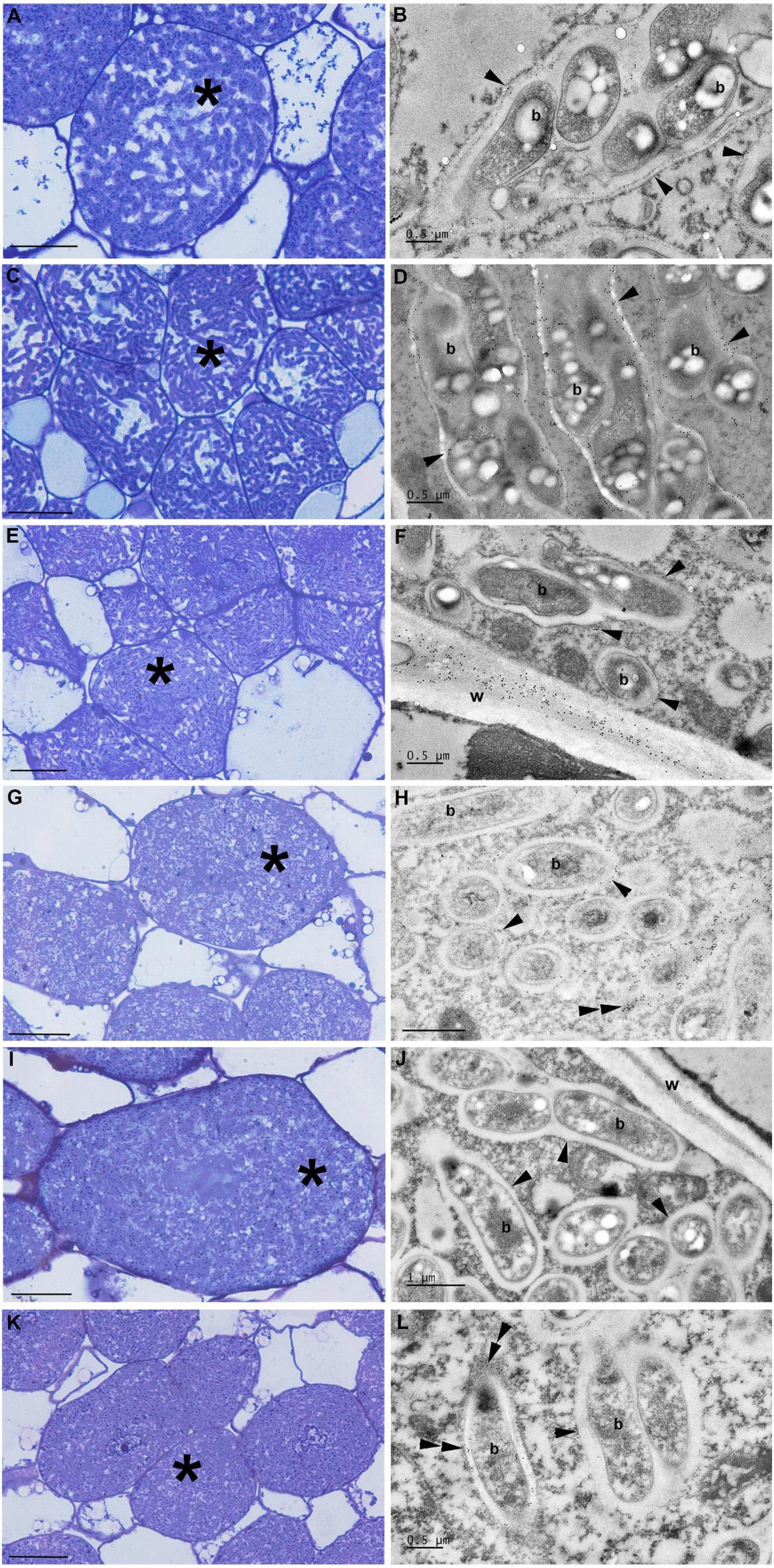


Fig. S2

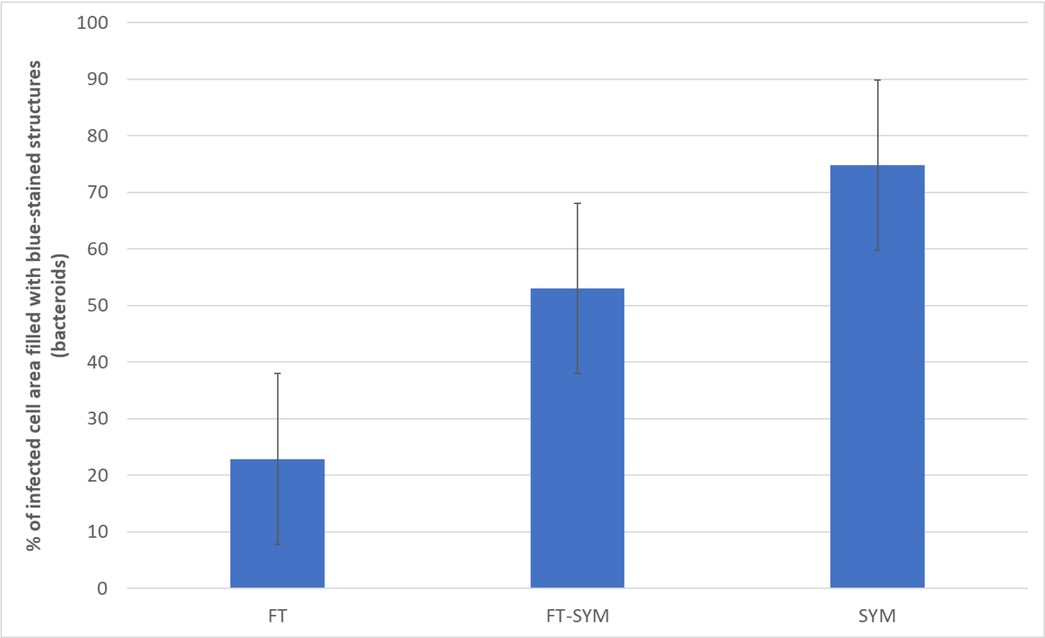


**FIG S3** Proportional (%) occupation of infected cells by symbiotic rhizobia estimated by counting pixels from light micrographs similar to those presented in Fig S2A and C for FT-type, Fig. S2E and G for FT-SYM-type, and Fig. S2I and K for SYM-type nodules. Data are presented as the mean proportion (± s.d.) of each infected cell profile filled with toluidine blue-stained structures representing bacteroids; 7-9 sections were examined per symbiosome type (FT, FT-SYM, SYM).

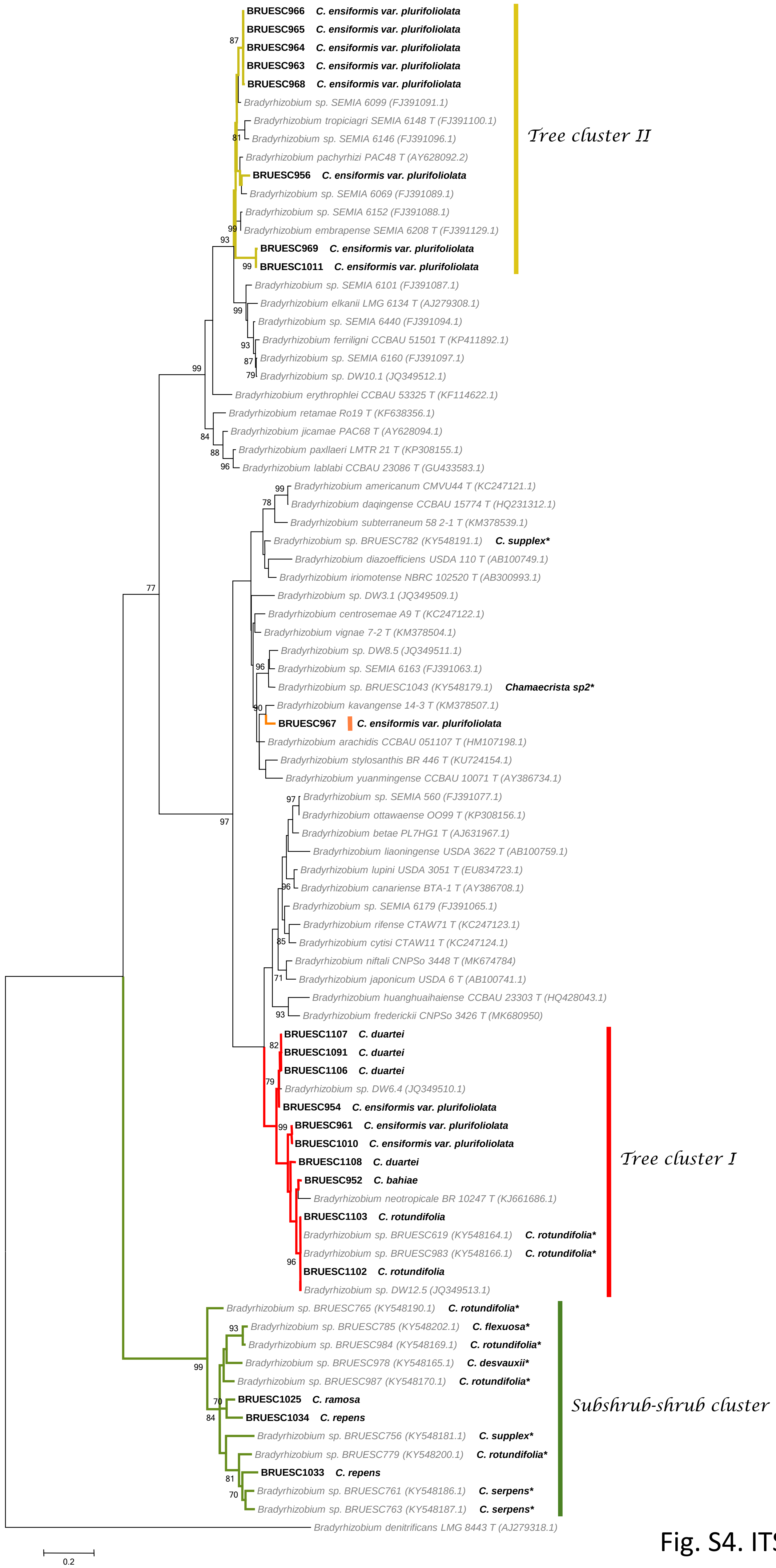
BRUESC966 C. ensiformis var. plurifoliolata
BRUESC965 C. ensiformis var. plurifoliolata
BRUESC964 C. ensiformis var. plurifoliolata
BRUESC963 C. ensiformis var. plurifoliolata
BRUESC968 C. ensiformis var. plurifoliolata
Bradyrhizobium sp. SEMIA 6099 (FJ391091.1)
Bradyrhizobium tropiciagri SEMIA 6148 T (FJ391100.1)
Bradyrhizobium sp. SEMIA 6146 (FJ391096.1)
Bradyrhizobium pachyrhizi PAC48 T (AY628092.2)
BRUESC956 C. ensiformis var. plurifoliolata
Bradyrhizobium sp. SEMIA 6069 (FJ391089.1)
Bradyrhizobium sp. SEMIA 6152 (FJ391088.1)
Bradyrhizobium embrapense SEMIA 6208 T (FJ391129.1)
BRUESC969 C. ensiformis var. plurifoliolata
BRUESC1011 C. ensiformis var. plurifoliolata
Tree cluster II
Bradyrhizobium sp. SEMIA 6101 (FJ391087.1)
Bradyrhizobium elkanii LMG 6134 T (AJ279308.1)
Bradyrhizobium sp. SEMIA 6440 (FJ391094.1)
Bradyrhizobium ferriligni CCBAU 51501 T (KP411892.1)
Bradyrhizobium sp. SEMIA 6160 (FJ391097.1)
Bradyrhizobium sp. DW10.1 (JQ349512.1)
Bradyrhizobium erythrophlei CCBAU 53325 T (KF114622.1)
Bradyrhizobium retamae Ro19 T (KF638356.1)
Bradyrhizobium jicamae PAC68 T (AY628094.1)
Bradyrhizobium paxllaeri LMTR 21 T (KP308155.1)
Bradyrhizobium lablabi CCBAU 23086 T (GU433583.1)
Bradyrhizobium americanum CMVU44 T (KC247121.1)
Bradyrhizobium daqingense CCBAU 15774 T (HQ231312.1)
Bradyrhizobium subterraneum 58 2-1 T (KM378539.1)
Bradyrhizobium sp. BRUESC782 (KY548191.1) C. supplex*
Bradyrhizobium diazoefficiens USDA 110 T (AB100749.1)
Bradyrhizobium iriomotense NBRC 102520 T (AB300993.1)
Bradyrhizobium sp. DW3.1 (JQ349509.1)
Bradyrhizobium centrosemae A9 T (KC247122.1)
Bradyrhizobium vignae 7-2 T (KM378504.1)
Bradyrhizobium sp. DW8.5 (JQ349511.1)
Bradyrhizobium sp. SEMIA 6163 (FJ391063.1)
Bradyrhizobium sp. BRUESC1043 (KY548179.1) Chamaecrista sp2*
Bradyrhizobium kavangense 14-3 T (KM378507.1)
BRUESC967 C. ensiformis var. plurifoliolata
Bradyrhizobium arachidis CCBAU 051107 T (HM107198.1)
Bradyrhizobium stylosanthis BR 446 T (KU724154.1)
Bradyrhizobium yuanmingense CCBAU 10071 T (AY386734.1)
Bradyrhizobium sp. SEMIA 560 (FJ391077.1)
Bradyrhizobium ottawaense OO99 T (KP308156.1)
Bradyrhizobium betae PL7HG1 T (AJ631967.1)
Bradyrhizobium liaoningense USDA 3622 T (AB100759.1)
Bradyrhizobium lupini USDA 3051 T (EU834723.1)
Bradyrhizobium canariense BTA-1 T (AY386708.1)
Bradyrhizobium sp. SEMIA 6179 (FJ391065.1)
Bradyrhizobium rifense CTAW71 T (KC247123.1)
Bradyrhizobium cytisi CTAW11 T (KC247124.1)
Bradyrhizobium niftali CNPSo 3448 T (MK674784)
Bradyrhizobium japonicum USDA 6 T (AB100741.1)
Bradyrhizobium huanghuaihaiense CCBAU 23303 T (HQ428043.1)
Bradyrhizobium frederickii CNPSo 3426 T (MK680950)
BRUESC1107 C. duartei
BRUESC1091 C. duartei
BRUESC1106 C. duartei
Bradyrhizobium sp. DW6.4 (JQ349510.1)
BRUESC954 C. ensiformis var. plurifoliolata
BRUESC961 C. ensiformis var. plurifoliolata
BRUESC1010 C. ensiformis var. plurifoliolata
BRUESC1108 C. duartei
BRUESC952 C. bahiae
Bradyrhizobium neotropicale BR 10247 T (KJ661686.1)
BRUESC1103 C. rotundifolia
Bradyrhizobium sp. BRUESC619 (KY548164.1) C. rotundifolia*
Bradyrhizobium sp. BRUESC983 (KY548166.1) C. rotundifolia*
BRUESC1102 C. rotundifolia
Bradyrhizobium sp. DW12.5 (JQ349513.1)
Tree cluster I
Bradyrhizobium sp. BRUESC765 (KY548190.1) C. rotundifolia*
Bradyrhizobium sp. BRUESC785 (KY548202.1) C. flexuosa*
Bradyrhizobium sp. BRUESC984 (KY548169.1) C. rotundifolia*
Bradyrhizobium sp. BRUESC978 (KY548165.1) C. desvauxii*
Bradyrhizobium sp. BRUESC987 (KY548170.1) C. rotundifolia*
BRUESC1025 C. ramosa
BRUESC1034 C. repens
Bradyrhizobium sp. BRUESC756 (KY548181.1) C. supplex*
Bradyrhizobium sp. BRUESC779 (KY548200.1) C. rotundifolia*
BRUESC1033 C. repens
Bradyrhizobium sp. BRUESC761 (KY548186.1) C. serpens*
Bradyrhizobium sp. BRUESC763 (KY548187.1) C. serpens*
Subshrub-shrub cluster
Bradyrhizobium denitrificans LMG 8443 T (AJ279318.1)
0.2

Fig. S4. ITS

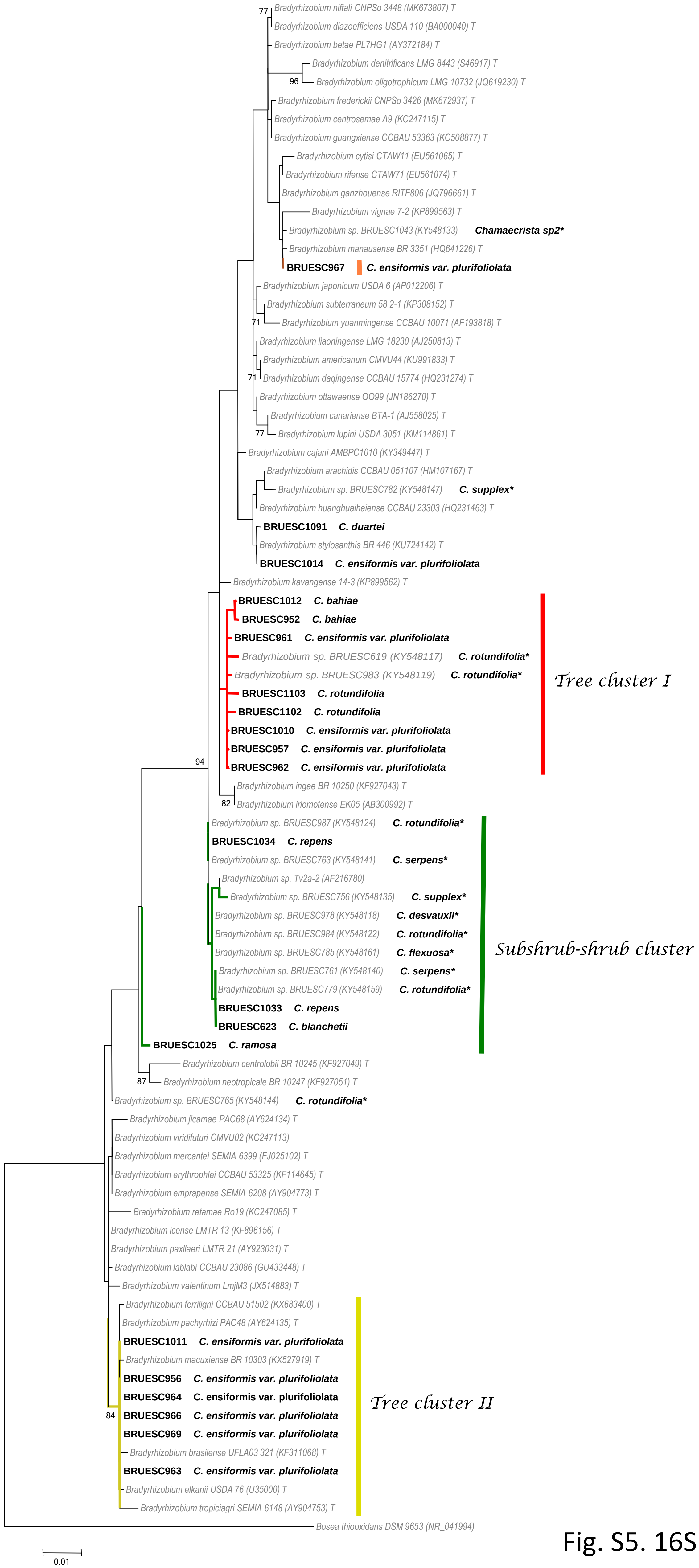


Fig. S5. 16S rDNA

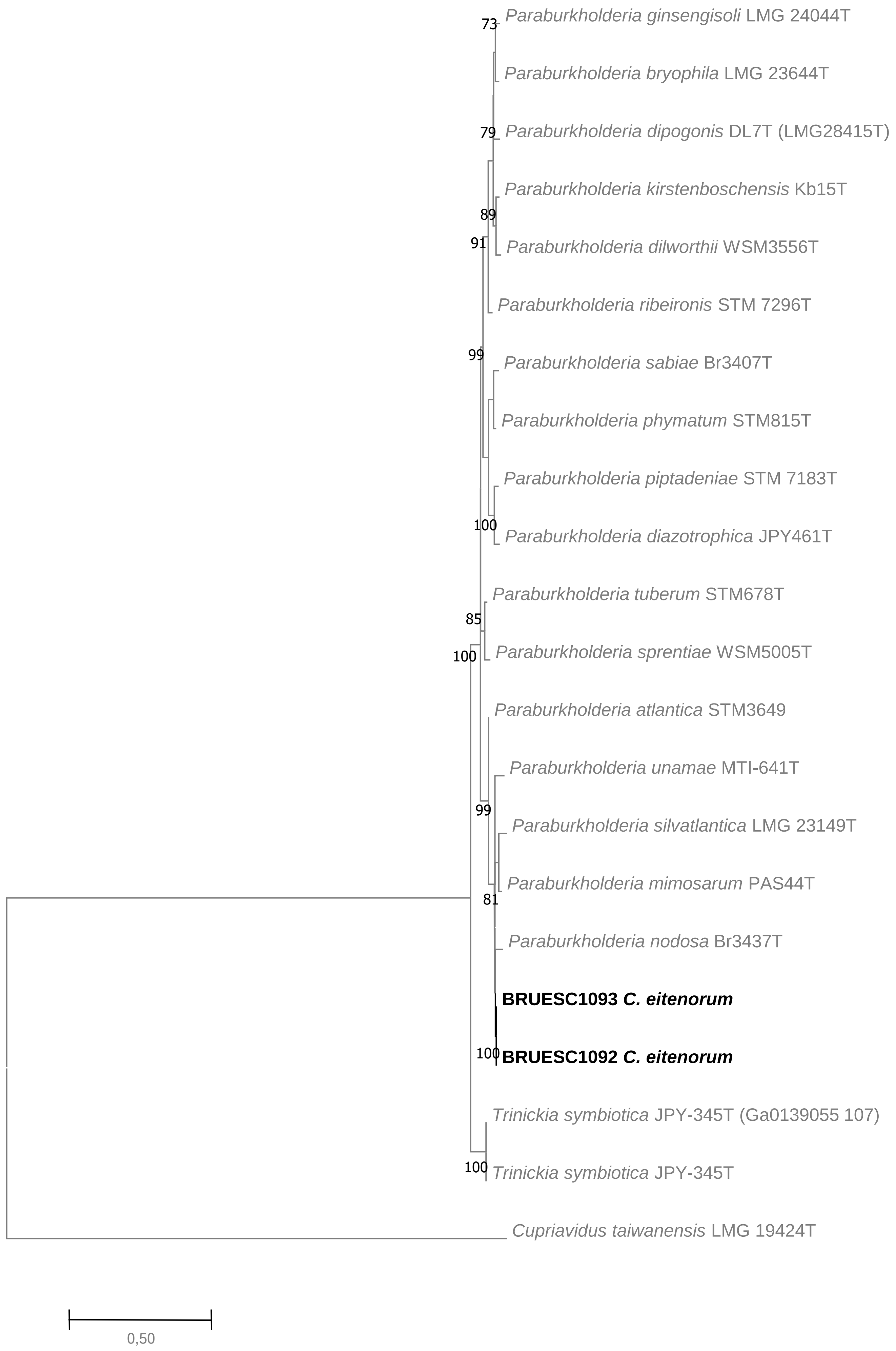


Fig. S6. 16S rDNA + *recA*

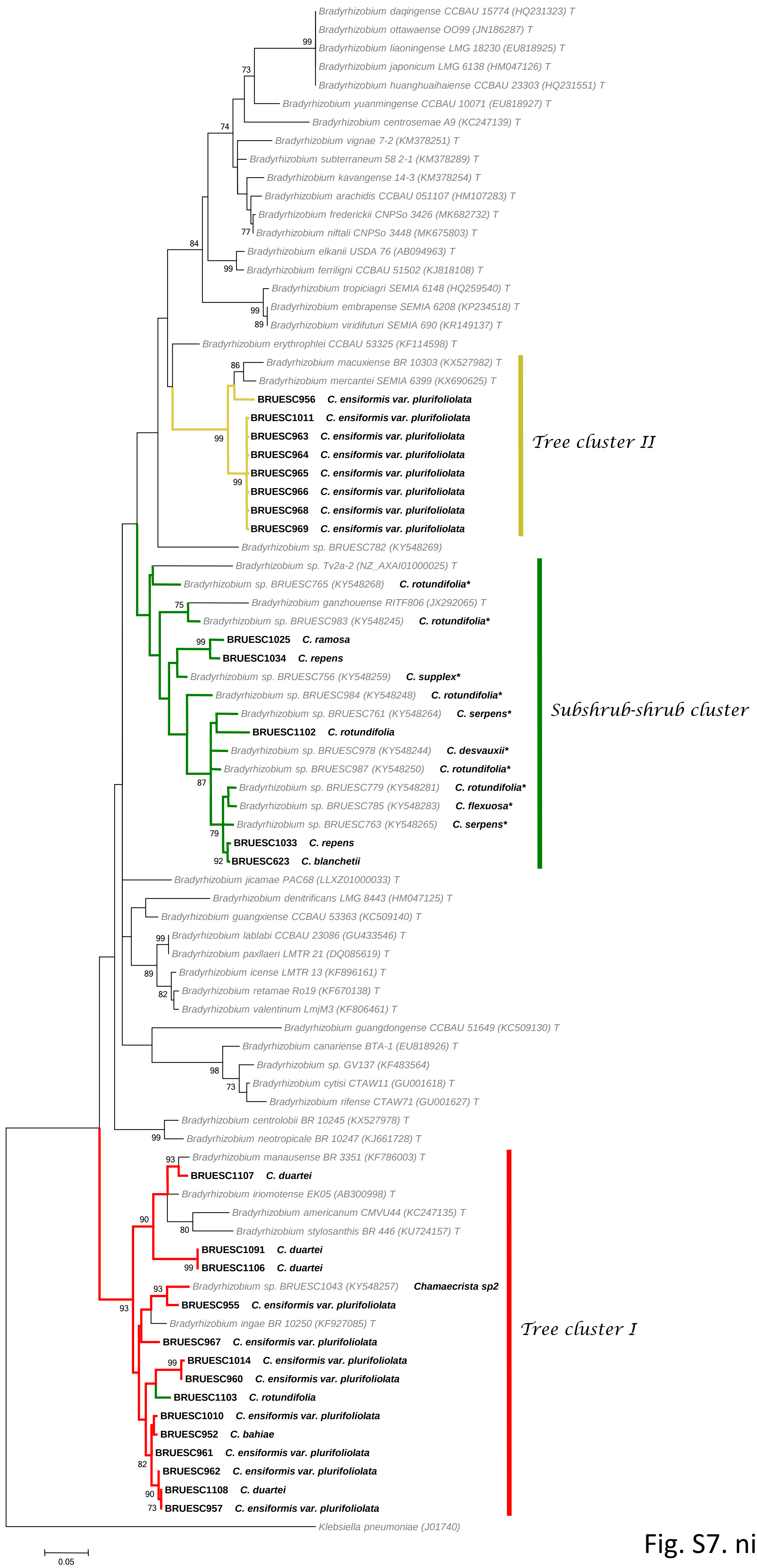

Bradyrhizobium daqingense CCBAU 15774 (HQ231323) T
Bradyrhizobium ottawaense OO99 (JN186287) T
Bradyrhizobium liaoningense LMG 18230 (EU818925) T
Bradyrhizobium japonicum LMG 6138 (HM047126) T
Bradyrhizobium huanghuaihaiense CCBAU 23303 (HQ231551) T
Bradyrhizobium yuanmingense CCBAU 10071 (EU818927) T
Bradyrhizobium centrosemae A9 (KC247139) T
Bradyrhizobium vignae 7-2 (KM378251) T
Bradyrhizobium subterraneum 58 2-1 (KM378289) T
Bradyrhizobium kavangense 14-3 (KM378254) T
Bradyrhizobium arachidis CCBAU 051107 (HM107283) T
Bradyrhizobium frederickii CNPSo 3426 (MK682732) T
Bradyrhizobium niftali CNPSo 3448 (MK675803) T
Bradyrhizobium elkanii USDA 76 (AB094963) T
Bradyrhizobium ferriligni CCBAU 51502 (KJ818108) T
Bradyrhizobium tropiciagri SEMIA 6148 (HQ259540) T
Bradyrhizobium embrapense SEMIA 6208 (KP234518) T
Bradyrhizobium viridifuturi SEMIA 690 (KR149137) T
Bradyrhizobium erythrophlei CCBAU 53325 (KF114598) T
Bradyrhizobium macuxiense BR 10303 (KX527982) T
Bradyrhizobium mercantei SEMIA 6399 (KX690625) T
BRUESC956 C. ensiformis var. plurifoliolata
BRUESC1011 C. ensiformis var. plurifoliolata
BRUESC963 C. ensiformis var. plurifoliolata
BRUESC964 C. ensiformis var. plurifoliolata
BRUESC965 C. ensiformis var. plurifoliolata
BRUESC966 C. ensiformis var. plurifoliolata
BRUESC968 C. ensiformis var. plurifoliolata
BRUESC969 C. ensiformis var. plurifoliolata
Tree cluster II
Bradyrhizobium sp. BRUESC782 (KY548269)
Bradyrhizobium sp. Tv2a-2 (NZ_AXAI01000025) T
Bradyrhizobium sp. BRUESC765 (KY548268) C. rotundifolia*
Bradyrhizobium ganzhouense RITF806 (JX292065) T
Bradyrhizobium sp. BRUESC983 (KY548245) C. rotundifolia*
BRUESC1025 C. ramosa
BRUESC1034 C. repens
Bradyrhizobium sp. BRUESC756 (KY548259) C. supplex*
Bradyrhizobium sp. BRUESC984 (KY548248) C. rotundifolia*
Bradyrhizobium sp. BRUESC761 (KY548264) C. serpens*
BRUESC1102 C. rotundifolia
Bradyrhizobium sp. BRUESC978 (KY548244) C. desvauxii*
Bradyrhizobium sp. BRUESC987 (KY548250) C. rotundifolia*
Bradyrhizobium sp. BRUESC779 (KY548281) C. rotundifolia*
Bradyrhizobium sp. BRUESC785 (KY548283) C. flexuosa*
Bradyrhizobium sp. BRUESC763 (KY548265) C. serpens*
BRUESC1033 C. repens
BRUESC623 C. blanchetii
Subshrub-shrub cluster
Bradyrhizobium jicamae PAC68 (LLXZ01000033) T
Bradyrhizobium denitrificans LMG 8443 (HM047125) T
Bradyrhizobium guangxiense CCBAU 53363 (KC509140) T
Bradyrhizobium lablabi CCBAU 23086 (GU433546) T
Bradyrhizobium paxllaeri LMTR 21 (DQ085619) T
Bradyrhizobium icense LMTR 13 (KF896161) T
Bradyrhizobium retamae Ro19 (KF670138) T
Bradyrhizobium valentinum LmjM3 (KF806461) T
Bradyrhizobium guangdongense CCBAU 51649 (KC509130) T
Bradyrhizobium canariense BTA-1 (EU818926) T
Bradyrhizobium sp. GV137 (KF483564)
Bradyrhizobium cytisi CTAW11 (GU001618) T
Bradyrhizobium rifense CTAW71 (GU001627) T
Bradyrhizobium centrolobii BR 10245 (KX527978) T
Bradyrhizobium neotropicale BR 10247 (KJ661728) T
Bradyrhizobium manausense BR 3351 (KF786003) T
BRUESC1107 C. duartei
Bradyrhizobium iriomotense EK05 (AB300998) T
Bradyrhizobium americanum CMVU44 (KC247135) T
Bradyrhizobium stylosanthis BR 446 (KU724157) T
BRUESC1091 C. duartei
BRUESC1106 C. duartei
Bradyrhizobium sp. BRUESC1043 (KY548257) Chamaecrista sp2
BRUESC955 C. ensiformis var. plurifoliolata
Bradyrhizobium ingae BR 10250 (KF927085) T
BRUESC967 C. ensiformis var. plurifoliolata
BRUESC1014 C. ensiformis var. plurifoliolata
BRUESC960 C. ensiformis var. plurifoliolata
BRUESC1103 C. rotundifolia
BRUESC1010 C. ensiformis var. plurifoliolata
BRUESC952 C. bahiae
BRUESC961 C. ensiformis var. plurifoliolata
BRUESC962 C. ensiformis var. plurifoliolata
BRUESC1108 C. duartei
BRUESC957 C. ensiformis var. plurifoliolata
Tree cluster I
Klebsiella pneumoniae (J01740)
0.05


Fig. S7. nifH

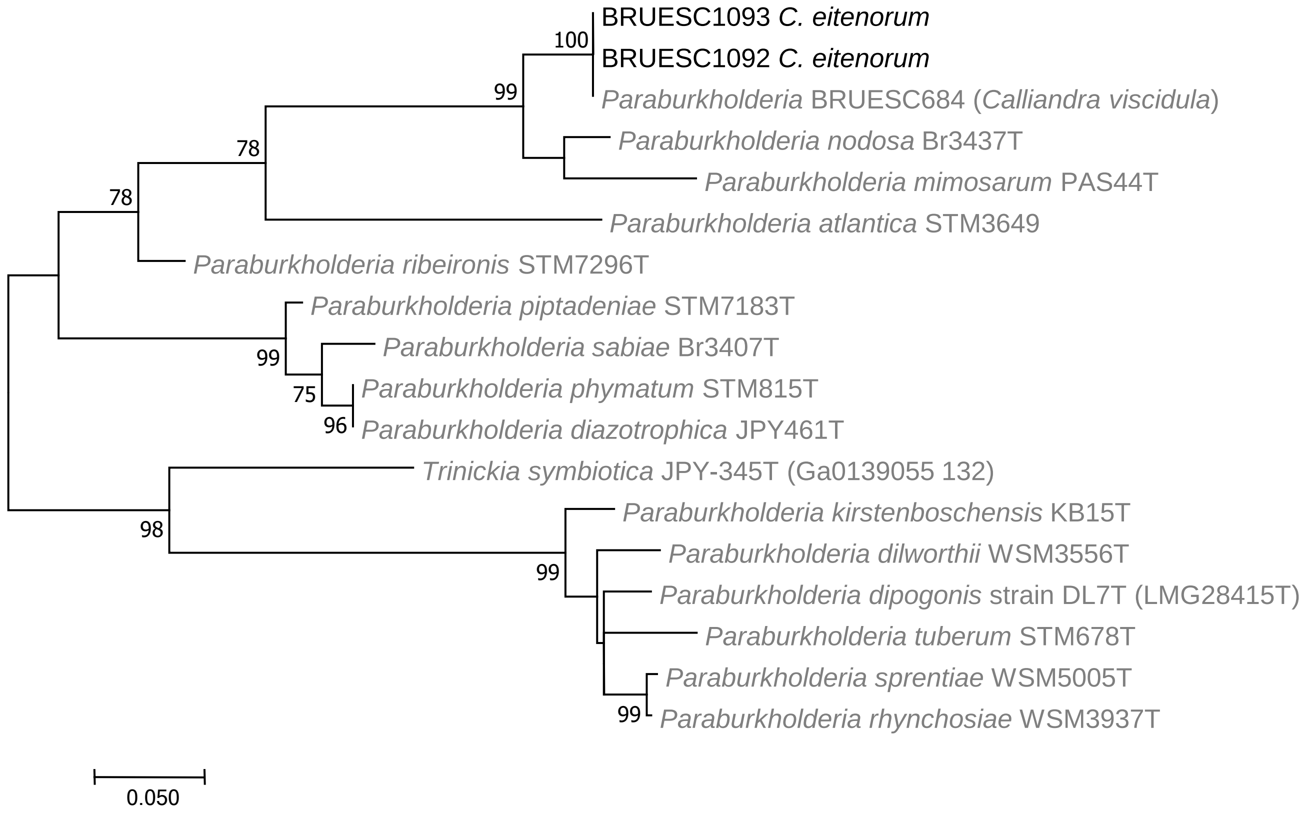
BRUESC1093 *C. eitenorum*
BRUESC1092 *C. eitenorum*
*Paraburkholderia* BRUESC684 (*Calliandra viscidula*)
*Paraburkholderia nodosa* Br3437T
*Paraburkholderia mimosarum* PAS44T
*Paraburkholderia atlantica* STM3649
*Paraburkholderia ribeironis* STM7296T
*Paraburkholderia piptadeniae* STM7183T
*Paraburkholderia sabiae* Br3407T
*Paraburkholderia phymatum* STM815T
*Paraburkholderia diazotrophica* JPY461T
*Trinickia symbiotica* JPY-345T (Ga0139055 132)
*Paraburkholderia kirstenboschensis* KB15T
*Paraburkholderia dilworthii* WSM3556T
*Paraburkholderia dipogonis* strain DL7T (LMG28415T)
*Paraburkholderia tuberum* STM678T
*Paraburkholderia sprentiae* WSM5005T
*Paraburkholderia rhynchosiae* WSM3937T
100
99
78
78
99
75
96
98
99
99
0.050

Fig. S8. *nodC*

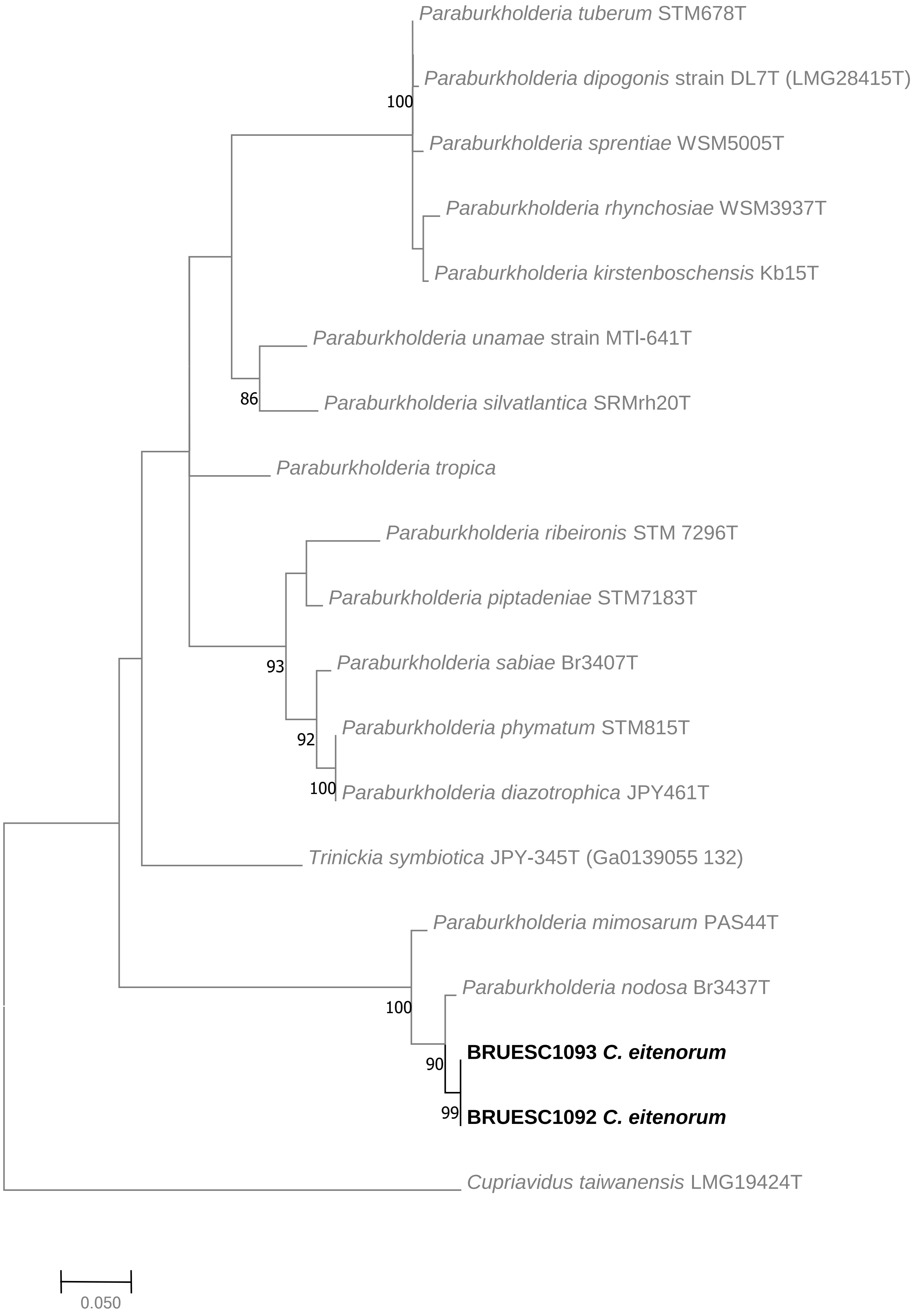
Paraburkholderia tuberum STM678T
Paraburkholderia dipogonis strain DL7T (LMG28415T)
100
Paraburkholderia sprentiae WSM5005T
Paraburkholderia rhynchosiae WSM3937T
Paraburkholderia kirstenboschensis Kb15T
Paraburkholderia unamae strain MTl-641T
86
Paraburkholderia silvatlantica SRMrh20T
Paraburkholderia tropica
Paraburkholderia ribeironis STM 7296T
Paraburkholderia piptadeniae STM7183T
93
Paraburkholderia sabiae Br3407T
92
Paraburkholderia phymatum STM815T
100
Paraburkholderia diazotrophica JPY461T
Trinickia symbiotica JPY-345T (Ga0139055 132)
Paraburkholderia mimosarum PAS44T
100
Paraburkholderia nodosa Br3437T
90
BRUESC1093 C. eitenorum
99
BRUESC1092 C. eitenorum
Cupriavidus taiwanensis LMG19424T
0.050

Fig. S9. nifH

**Table S1.** Accession numbers of aerial parts of *Chamaecrista* species sampled in Bahia (BA), Brazil and deposited in the Herbarium of the Universidade Estadual de Santa Cruz (UESC). Also included is information about growth habit, vegetation type and locality where the *Chamaecrista* species were sampled in BA specifically for this study.

| **Species** | **Herbarium voucher ID** | **Habit** | **Vegetation type** | **Locality** | **Altitude (m)** | **Latitude** | **Longitude** |
|---|---|---|---|---|---|---|---|
| *C. bahiae* (H.S.Irwin) H.S.Irwin & Barneby | 17198 | Tree | Rf | Prado | 35 | 17º15’07’’S | 39°13'10”W |
| *C. duartei* (H.S.Irwin) H.S.Irwin & Barneby | 18541 | Tree | Rf | Ilhéus | 26 | 14º47’50”S | 39°10'18”W |
| C*. ensiformis* var. plurifoliolata (Hoehne) H.S.Irwin & Barneby | 17204 | Tree | Rf | Ibicaraí | 102 | 14º54’22”S | 39°36'44”W |
| *C. eitenorum* H.S.Irwin & Barneby | 21236 | Tree | SDTF | Lençóis | 450 | 12°33’58”S | 41°23’49”W |
| *C. blanchetii* (Benth.) Conc. et al. | 19930 | Treelet to shrub | *Cr* | Jacobina | 658 | 11°11’40”S | 40°30’36”W |
| *C.* × *blanchetiformis* Conc. et al | 19913 | Treelet to shrub | *Cr* | Mucugê | 888 | 12°59’01.8”S | 41°21’31.6”W |
| *C. brachystachya* (Benth.) Conc. et al. | 19918 | Treelet to shrub | *Cr* | Mucugê | 815 | 12°57’01.8”S | 41°16’43”W |
| *C. confertiformis* (H.S.Irwin & Barneby) Conc. et al. | 19914 | Treelet to shrub | *Cr* | Mucugê | 980 | 12°59’59.6”S | 41°23’23.2”W |
| *C. zygophylloides* (Taub.) H.S.Irwin & Barneby | 21234 | Shrub | *Cr* | Lençóis | 487 | 12°33’38.3”S | 41°24’14.1”W |
| *C. belemii* (H.S.Irwin & Barneby) H.S.Irwin & Barneby | 21237 | Shrub | *Cr* | Morro do Chapéu | 927 | 11°29’26”S | 41°19’54.1”W |

| | | | | | | | |
|---|---|---|---|---|---|---|---|
| *C. arrojadoana* (Harms) Rando | 19935 | Shrub | *Cr* | Lençóis | 701 | 12°27’43.9”S | 41°24’53.7”W |
| *C. repens* (Vogel) H.S.Irwin & Barneby | 16298 | Shrub | *Ca* | Morro do Chapéu | 765 | 10º21’47”S | 40º12’06”W |
| *C. repens (*Vogel) H.S.Irwin & Barneby | 19934 | Shrub | *Cr* | Lençóis | 450 | 12º33’52”S | 41º23’42.7”W |
| *C. repens* (Vogel) H.S.Irwin & Barneby | 21235 | Shrub | *Cr* | Lençóis | 454 | 12º34’14.5”S | 41º23’26.5”W |
| *C. ramosa* (Vogel) H.S.Irwin & Barneby | 19932 | Shrub to subshrub | *Cr* | Lençóis | 399 | 13º48’53.9”S | 42º40’06”W |
| *C. desvauxii* (Collad.) Killip | 17196 | Shrub to subshrub | *Cr*, *Ca* and *R* | Remanso | 417 | 09°45’34.3”S | 42°17’54.1”W |
| *C. desvauxii* (Collad.) Killip | 17196 | Shrub to subshrub | *Cr*, *Ca* and *R* | Remanso | 417 | 09°45’34.3”S | 42°17’54.1”W |
| *C. pascuorum* (Benth.) H.S.Irwin & Barneby | 19937 | Subshrub | *Cr* and *Ca* | Caetité | 889 | 14°47’47.4”S | 39°10’26.7”W |
| *C. rotundifolia* (Pers.) Greene | 17419 | Subshrub | *Ca* | Pilão Arcado | 424 | 10º08’06.4”S | 42º54’57.3”W |
| *C. supplex* (Mart. ex Benth.) Britton & Rose ex Britton & Killip | 17417 | Subshrub | *Ca* | Remanso | 368 | 09°36’16.2”S | 42°06’05.7”W |
| *C. serpens* (L.) Grenee | 17427 | Subshrub | *Ca* | Remanso | 412 | 09°45’33.1”S | 42°17’54.0”W |
| *C. flexuosa* (L.) Grenee | 21438 | Subshrub | *Cr* | Morro do Chapéu | 893 | 11°37’35.2”S | 41°00’06.3”W |

Rf = rainforest; Cr = campo rupestre; Ca = caatinga; R = restinga; SDTF = seasonally dry tropical forest.

**Table S3.** Voucher information and GenBank accession numbers of the nuclear ITS and plastidial *trnL-trnF* loci sequences included in the phylogeny of *Chamaecrista* (Fig. 8). New sequences are marked with an asterisk (*).

| ***Species*** | **Voucher** | **ITS** | **trnL-F** |
|---|---|---|---|
| *Cassia grandis* L. | L.P. Queiroz 2878 (HUEFS) | FJ009820 | FJ009875 |
| *Cassia javanica* L. | L.P. Queiroz 11039 (HUEFS) | FJ009821 | FJ009876 |
| *Senna alata* (L.) Roxb. | Bruneau 1076 (K) and D.J. Gomes 42 (HUEFS) | KX372780 | MW075531* |
| *Senna gardneri* (Benth.) H.S. Irwin & Barneby | L.P. Queiroz 7860 (HUEFS) | FJ009822 | FJ009877 |
| *Chamaecrista absus* (L.) H.S. Irwin & Barneby var. *absus* | Conceição 1056 (HUEFS) | FJ009832 | FJ009886 |
| *Chamaecrista adiantifolia* (Spruce ex Benth.) H.S. Irwin & Barneby | J.G. Rando 1197 (HUEFS) | MW075532* | MW075526* |
| *Chamaecrista altoana* (H.S. Irwin & Barneby) H.S. Irwin & Barneby | M.J. Silva et al. 6465 (UFG) | MH806439 | MH828377 |
| *Chamaecrista amorimii* Barneby | Conceição 795 (HUEFS) | FJ009823 | FJ009878 |
| *Chamaecrista anamariae* Conc., L.P. Queiroz & G.P. Lewis | Conceição 787 (HUEFS) | FJ009826 | FJ009881 |
| *Chamaecrista aristata* (Benth.) H.S. Irwin & Barneby | J.G. Rando 976 (SPF) | KP967070 | KP966899 |
| *Chamaecrista arrojadoana* (Harms) Rando | J.G. Rando 1011 (HUEFS) | KP967093 | KP966920 |
| *Chamaecrista azulana* (H.S. Irwin & Barneby) H.S. Irwin & Barneby | A.O. Souza 1116 (UFG) | MH835355 | MH828378 |
| *Chamaecrista bahiae* (H.S. Irwin) H.S. Irwin & Barneby | J.G. Rando JG 1213 (SPF) | x | MW075525* |
| *Chamaecrista belemii* (H.S. Irwin & Barneby) H.S. Irwin & Barneby var. *belemii* | L.P. Queiroz | DQ787389 | FJ009880 |

| | | | |
|---|---|---|---|
| | 9151 (HUEFS) | | |
| *Chamaecrista beladona* Silva & Souza | M.J. Silva 5993 (UFG | MH828379 | MH828399 |
| *Chamaecrista benthamiana* (Harms) H.S. Irwin & Barneby | A.O. Souza & al. 1576 (UFG) | MH835357 | MH828380 |
| *Chamaecrista blanchetii* Conc., L.P. Queiroz & G.P. Lewis | Andrade 607 (HUEFS) | FJ009846 | FJ009900 |
| *Chamaecrista botryoides* Conc., L.P. Queiroz & G.P. Lewis | Conceição 541 (HUEFS) | FJ009836 | FJ009890 |
| *Chamaecrista brachyblepharis* (Harms) H.S. Irwin & Barneby | M.J. Silva 4472 (UFG) | MH835358 | MH828381 |
| *Chamaecrista brachyrachis* (Harms) H.S. Irwin & Barneby | A.O. Souza & al. 1179 (UFG) | MH835359 | MH828382 |
| *Chamaecrista brachystachya* (Benth.) Conc., L.P. Queiroz & G.P. Lewis | Conceição 713 (HUEFS) | FJ009847 | FJ009901 |
| *Chamaecrista brevicalyx* (Benth.) H.S. Irwin & Barneby var. *brevicalyx* | A.O. Souza & al. 1245 (UFG) | MH835360 | MH828383 |
| *Chamaecrista burchellii* (Benth.) H.S. Irwin & Barneby | J.G. Rando 1092 (HUEFS, SPF) | KP967073 | KP966900 |
| *Chamaecrista calycioides* (DC. ex Collad.) Greene | Queiroz 11 (HUEFS) | FJ009863 | FJ009917 |
| *Chamaecrista campestris* H.S. Irwin & Barneby 1 | L.P. Queiroz 10440 (HUEFS) | FJ009829 | FJ009883 |
| *Chamaecrista campestris* H.S. Irwin & Barneby 2 | A.O. Souza & al. 1596 (UFG) | MH835361 | MH828384 |
| *Chamaecrista campicola* (Harms) H.S. Irwin & Barneby | A.O. Souza 893 (UFG) | MH835362 | MH828385 |
| *Chamaecrista cardiostegia* H.S. Irwin & Barneby | J.G. Rando 1125 (SPF) | KP967074 | KP966901 |
| *Chamaecrista caribaea* var. *lucayana* (Britton) H.S. Irwin & Barneby | J.G. Rando 963 (NY, SPF) | KP967075 | KP966902 |
| *Chamaecrista cathartica* (Mart.) H.S. Irwin & Barneby | Conceição 789 (HUEFS) | FJ009841 | FJ009895 |
| *Chamaecrista* cf. *isidorea* (Benth.) H.S. Irwin & Barneby | M.J. Silva 6059 (UFG) | MH835363 | MH828386 |

| | | | |
|---|---|---|---|
| *Chamaecrista choriophylla* (Vogel) H.S. Irwin & Barneby | J.G. Rando 1034 (HUEFS) | KP967076 | KP966904 |
| *Chamaecrista ciliolata* (Benth.) H.S. Irwin & Barneby var*. ciliolata* | J.G. Rando 1115 (HUEFS, SPF) | MH835364 | MH828387 |
| *Chamaecrista ciliolata* var*. pulchella* (H.S. Irwin & Barneby) H.S. Irwin & Barneby | A.O. Souza 1423 (UFG) | MH835365 | MH828388 |
| *Chamaecrista cinerascens* (Vogel) H.S. Irwin & Barneby | J.G. Rando 661 (SPF) | KP967077 | KP966960 |
| *Chamaecrista cipoana* (H.S. Irwin & Barneby) H.S. Irwin & Barneby | A.O. Souza 1414 (UFG) | MH835366 | MH828389 |
| *Chamaecrista clausseni* (Benth.) H.S. Irwin & Barneby*i* var*. claussenii* | A.O. Souza 979 (UFG) | MH835367 | MH828390 |
| *Chamaecrista confertiformis* (H.S. Irwin & Barneby) Conc., L.P. Queiroz & G.P. Lewis | Costa 132 (HUEFS) | FJ009848 | FJ009902 |
| *Chamaecrista coradinii* H.S. Irwin & Barneby | L.L.C. Antunes 1579 (UFG) | MH835368 | MH828391 |
| *Chamaecrista coriacea* (Bong. ex Benth.) H.S. Irwin & Barneby | Conceição 869 (HUEFS) | FJ009843 | FJ009897 |
| *Chamaecrista cytisoides* (DC. ex Collad.) H.S. Irwin & Barneby | Conceição 870 (HUEFS) | FJ009844 | FJ009898 |
| *Chamaecrista dalbergiifolia* (Benth.) H.S. Irwin & Barneby | L.P. Queiroz 10318 (HUEFS) | FJ009837 | FJ009891 |
| *Chamaecrista dawsonii* (R.S. Cowan) H.S. Irwin & Barneby | M.J. Silva 4474 (UFG) | MH835369 | MH828392 |
| *Chamaecrista decora* (H.S. Irwin & Barneby) Conc., L.P. Queiroz & G.P. Lewis | Conceição 810 (HUEFS) | FJ009849 | FJ009903 |
| *Chamaecrista decumbens* (Benth.) H.S. Irwin & Barneby | A.O. Souza 802 (UFG) | MH835370 | MH828393 |
| *Chamaecrista densifolia* (Benth.) H.S. Irwin & Barneby | A.O. Souza 1159 (UFG) | MH835371 | MH828394 |
| *Chamaecrista depauperata* Conc., L.P. Queiroz & G.P. Lewis | Conceição 863 (HUEFS) | FJ009850 | FJ009904 |
| *Chamaecrista desvauxii* (Collad.) Killip var*. desvauxii* | L.P. Queiroz 10453 (HUEFS) | FJ009864 | FJ009918 |

| | | | |
|---|---|---|---|
| *Chamaecrista desvauxii* var. *langsdorfii* (Kunth ex Vogel) H.S. Irwin & Barneby | Conceição 674 (HUEFS) | FJ009866 | FJ009920 |
| *Chamaecrista desvauxii* var. *latistipula* (Benth.) G.P. Lewis | Conceição 912 (HUEFS) | FJ009867 | FJ009921 |
| *Chamaecrista desvauxii* var. *mollissima* (Benth.) H.S. Irwin & Barneby | Santos 356 (HUEFS) | FJ009865 | FJ009919 |
| *Chamaecrista diphylla* (L.) Greene | L.P. Queiroz 10269 (HUEFS) | FJ009868 | FJ009922 |
| *Chamaecrista distichoclada* (Mart. ex Benth.) H.S. Irwin & Barneby | J.G. Rando 1230 (HUEFS, SPF) | KP967078 | KP966961 |
| *Chamaecrista eitenorum* (H.S. Irwin & Barneby) H.S. Irwin & Barneby | Cardoso 3483 (HUEFS) | MW075533* | MW075529* |
| *Chamaecrista elata* A.O. Souza & M.J. Silva | M.J. Silva 6187 (UFG) | MH835372 | MH828395 |
| *Chamaecrista ensiformis* var. *plurifoliolata* (Hoehne) H.S. Irwin & Barneby | L.P. Queiroz 13879 (HUEFS) | MW075534* | MW075530* |
| *Chamaecrista fagonioides* var. *macrocalyx* (H.S. Irwin & Barneby) H.S. Irwin & Barneby | A.O. Souza 910 (UFG) | MH835373 | MH828396 |
| *Chamaecrista fasciculata* (Michx.) Greene | | QANZ01038945.1 | QANZ01049895.1 |
| *Chamaecrista feliciana* (H.S. Irwin & Barneby) H.S. Irwin & Barneby | A.O. Souza 1500 (UFG) | MH835374 | MH828397 |
| *Chamaecrista filicifolia* (Mart. ex Benth.) H.S. Irwin & Barneby | A.O. Souza 781 (UFG) | MH835375 | MH828398 |
| *Chamaecrista flexuosa* (L.) Greene | Giulietti 2344 (HUEFS) | FJ009858 | FJ009912 |
| *Chamaecrista floribunda* M.J. Silva & A.O. Souza | A.O. Souza 1288 (UFG) | MH835376 | MH828399 |
| *Chamaecrista glaucofilix* (H.S. Irwin & Barneby) H.S. Irwin & Barneby 1 | Conceição 861, HUEFS | FJ009834 | FJ009888 |
| *Chamaecrista glaucofilix* (H.S. Irwin & Barneby) H.S. Irwin & Barneby 2 | A.O. Souza 1550 (UFG) | MH835377 | MH828400 |
| *Chamaecrista gymnothyrsa* (H.S. Irwin & Barneby) H.S. Irwin & Barneby | R.C. Sodre 1330 (UFG) | MH835378 | MH828401 |

| | | | |
|---|---|---|---|
| *Chamaecrista hispidula* (Vahl) H.S. Irwin & Barneby | Conceição 914 (HUEFS) | FJ009833 | FJ009887 |
| *Chamaecrista irwiniana* A.O. Souza & M.J. Silva | A.O. Souza 605 (UFG) | MH835380 | MH828403 |
| *Chamaecrista jacobinea* (Benth.) H.S. Irwin & Barneby | Andrade 610 (HUEFS) | FJ009827 | FJ009882 |
| *Chamaecrista lagotois* H.S. Irwin & Barneby | J.G. Rando 1029 (HUEFS) | KP967079 | KP966963 |
| *Chamaecrista latifolia* (Benth.) Rando | J.G. Rando 1024 (HUEFS) | KP967081 | KP966909 |
| *Chamaecrista lineata* (Sw.) Greene | J.G. Rando 958 (SPF) | KP967085 | KP966967 |
| *Chamaecrista macedoi* (H.S. Irwin & Barneby) H.S. Irwin & Barneby | M.J. Silva 6073 (UFG) | MH835381 | MH828404 |
| *Chamaecrista mollicaulis* (Harms) H.S. Irwin & Barneby | M.J. Silva 5693 (UFG) | MH835382 | MH828405 |
| *Chamaecrista mucronata* (Spreng.) H.S. Irwin & Barneby | J.G. Rando 879 (SPF) | KP967086 | KP966913 |
| *Chamaecrista multipennis* (H.S. Irwin & Barneby) H.S. Irwin & Barneby | A.O. Souza 1411 (UFG) | MH835383 | MH828406 |
| *Chamaecrista multiseta* (Benth.) H.S. Irwin & Barneby | A.O. Souza & al. 1763 (UFG) | MH835385 | MH828408 |
| *Chamaecrista nanodes* (H.S. Irwin & Barneby) H.S. Irwin & Barneby | A.O. Souza 1763 (UFG) | MH835385 | MH828408 |
| *Chamaecrista neesiana* var. *laxiracemosa* (Harms) H.S. Irwin & Barneby | M.J. Silva 5183 (UFG) | MH835386 | MH828409 |
| *Chamaecrista negrensis* (H.S. Irwin) H.S. Irwin & Barneby | Cardoso 3395 (HUEFS) | MW075535* | MW075528* |
| *Chamaecrista nictitans* subsp*. brachypoda* (Benth.) H.S. Irwin & Barneby | L.P. Queiroz 10335 (HUEFS) | FJ009855 | FJ009909 |
| *Chamaecrista nictitans* subsp*. disadena* (Steud.) H.S. Irwin & Barneby var*. disadena* | Conceição 790 (HUEFS) | FJ009852 | FJ009906 |
| *Chamaecrista nictitans* subsp*. patellaria* var*. ramosa* (Vogel) H.S. Irwin & Barneby | L.P. Queiroz 10406 (HUEFS) | FJ009853 | FJ009907 |

| | | | |
|---|---|---|---|
| *Chamaecrista nummulariifolia* (Benth.) H.S. Irwin & Barneby | A.O. Souza 790 (UFG) | MH835387 | MH828410 |
| *Chamaecrista obolaria* A.O. Souza & M.J. Silva | A.O. Souza 864 (UFG) | MH835388 | MH828411 |
| *Chamaecrista olesiphylla* (Vogel) H.S. Irwin & Barneby | J.G. Rando 1147 (SPF) | KP967089 | KP966916 |
| *Chamaecrista onusta* H.S. Irwin & Barneby | Conceição 800 (HUEFS) | FJ009824 | FJ009879 |
| *Chamaecrista pachyclada* (Harms) H.S. Irwin & Barneby | A.O. Souza 557 (UFG) | MH835389 | MH828412 |
| *Chamaecrista pascuorum* (Mart. ex Benth.) H.S. Irwin & Barneby | L.P. Queiroz 9169 (HUEFS) | FJ009851 | FJ009905 |
| *Chamaecrista philippi* (H.S. Irwin & Barneby) H.S. Irwin & Barneby | Giulietti 2245 (HUEFS) | FJ009838 | FJ009892 |
| *Chamaecrista pilosa* (L.) Greene | L.P. Queiroz 10221 (HUEFS) | FJ009856 | FJ009910 |
| *Chamaecrista planaltoana* (Harms) H.S. Irwin & Barneby | A.O. Souza 1355 (UFG) | MH835390 | MH828413 |
| *Chamaecrista polita* (H.S. Irwin & Barneby) H.S. Irwin & Barneby | A.O. Souza 352 (UFG) | MH835391 | MH828414 |
| *Chamaecrista potentilla* (Mart. ex Benth.) H.S. Irwin & Barneby | J.G. Rando 1139 (HUEFS, SPF) | KP967096 | KP966923 |
| *Chamaecrista pumila* (Lam.) V. Singh | | MH768080 | KU551117 |
| *Chamaecrista ramosa* var. *parvifoliola* (H.S. Irwin) H.S. Irwin & Barneby | Barbosa 787 (HUEFS) | KR134043 | x |
| *Chamaecrista ramosa* (Vogel) H.S. Irwin & Barneby var. *ramosa* | Barbosa 766 (HUEFS) | KR134010 | x |
| *Chamaecrista repens* (Vogel) H.S. Irwin & Barneby | Giulietti 2325 (HUEFS) | KP967098 | KP966925 |
| *Chamaecrista roraimae* (Benth.) Gleason | J.G. Rando 1154 (SPF) | KP967099 | KP966927 |
| *Chamaecrista rossicorum* (H.S. Irwin & Barneby) Rando | J.G. Rando 1045 (HUEFS, SPF) | KP967101 | KP966931 |
| *Chamaecrista rotundata* var. *interstes* H.S. Irwin & Barneby | J.G. Rando 1145 (SPF) | KP967107 | KP966938 |

| | | | |
|---|---|---|---|
| *Chamaecrista rotundata* (Vogel) H.S. Irwin & Barneby var. *rotundata* | J.G. Rando 925 (K, SPF) | KP967104 | KP966934 |
| *Chamaecrista rotundifolia* var*. grandiflora* (Benth.) H.S. Irwin & Barneby | Costa 128 (HUEFS) | FJ009857 | FJ009911 |
| *Chamaecrista rupestrium* H.S. Irwin & Barneby 1 | Santos 390 (HUEFS) | FJ009835 | FJ009889 |
| *Chamaecrista rupestrium* H.S. Irwin & Barneby 2 | A.O. Souza 1234 (UFG) | MH835392 | MH828415 |
| *Chamaecrista scabra* (Pohl ex Benth.) H.S. Irwin & Barneby | A.O. Souza 1456 (UFG) | MH835393 | MH828416 |
| *Chamaecrista serpens* (L.) Greene | L.P. Queiroz 10041 (HUEFS) | x | MW075524* |
| *Chamaecrista setosa var detonsa* (Benth.) H.S. Irwin & Barneby 1 | L.P. Queiroz 10460 (HUEFS) | FJ009842 | FJ009896 |
| *Chamaecrista setosa var detonsa* (Benth.) H.S. Irwin & Barneby 2 | A.O. Souza 405 (UFG) | MH835394 | MH828417 |
| *Chamaecrista simplifacta* H.S. Irwin & Barneby | J.G. Rando 1148 (SPF) | KP967110 | KP966943 |
| *Chamaecrista sincorana* (Harms) H.S. Irwin & Barneby | A.O. Souza 1246 (UFG) | MH835395 | MH828418 |
| *Chamaecrista sparsifolia* A.O. Souza & M.J. Silva | A.O. Souza 1050 (UFG) | MH835396 | MH828419 |
| *Chamaecrista speciosa* Conc., L.P. Queiroz & G.P. Lewis | Conceição 546 (HUEFS) | FJ009839 | FJ009893 |
| *Chamaecrista strictula* (H.S. Irwin & Barneby) H.S. Irwin & Barneby | A.O. Souza 1324 (UFG) | MH835397 | MH828420 |
| *Chamaecrista supplex* (Mart. ex Benth.) Britton & Rose ex Britton & Killip | L.P. Queiroz 10217 (HUEFS) | FJ009869 | FJ009923 |
| *Chamaecrista swainsonii* (Benth.) H.S. Irwin & Barneby | L.P. Queiroz 12314 (HUEFS) | KP967111 | KP966944 |
| *Chamaecrista tenuicaulis* A.O. Souza & M.J. Silva | A.O. Souza 1256 (UFG) | MH835398 | MH828421 |
| *Chamaecrista tragacanthoides* var*. rasa* H.S. Irwin & Barneby | J.G. Rando 1005 (SPF) | KP967113 | KP967113 |
| *Chamaecrista tragacanthoides* (Mart. ex | Pirani 6334 (SPF) | KP967114 | KP966948 |

| | | | |
|---|---|---|---|
| Benth.) H.S. Irwin & Barneby var. *tragacanthoides* | | | |
| *Chamaecrista ulmea* H.S. Irwin & Barneby | Santos 650 (SPF) | KP967115 | KP966949 |
| *Chamaecrista unijuga* (Benth.) Conc., L.P. Queiroz & G.P. Lewis | Conceição 694 (HUEFS) | FJ009845 | FJ009899 |
| *Chamaecrista urophyllidia* (H.S. Irwin & Barneby) H.S. Irwin & Barneby | Harley 54656 (HUEFS) | FJ009840 | FJ009894 |
| *Chamaecrista venulosa* (Benth.) H.S. Irwin & Barneby | J.G. Rando 1015 (HUEFS) | KP967116 | KP966950 |
| *Chamaecrista viscosa* var. *paraguariensis* (Chodat & Hassl.) H.S. Irwin & Barneby | A.O. Souza 1641 (UFG) | MH835399 | MH828422 |
